# Baby Open Brains: An Open-Source Repository of Infant Brain Segmentations

**DOI:** 10.1101/2024.10.02.616147

**Authors:** Eric Feczko, Sally M Stoyell, Lucille A. Moore, Dimitrios Alexopoulos, Maria Bagonis, Kenneth Barrett, Brad Bower, Addison Cavender, Taylor A. Chamberlain, Greg Conan, Trevor KM Day, Dhruman Goradia, Alice Graham, Lucas Heisler-Roman, Timothy J. Hendrickson, Audrey Houghton, Omid Kardan, Elizabeth A Kiffmeyer, Erik G Lee, Jacob T. Lundquist, Carina Lucena, Tabitha Martin, Anurima Mummaneni, Mollie Myricks, Pranav Narnur, Anders J. Perrone, Paul Reiners, Amanda R. Rueter, Hteemoo Saw, Martin Styner, Sooyeon Sung, Barry Tiklasky, Jessica L Wisnowski, Essa Yacoub, Brett Zimmermann, Christopher D. Smyser, Monica D. Rosenberg, Damien A. Fair, Jed T. Elison

**Author notes:** Co-first authors (Eric Feczko, Sally M Stoyell).

## Abstract

Reproducibility of neuroimaging research on infant brain development remains limited due to highly variable protocols and processing approaches. Progress towards reproducible pipelines is limited by a lack of benchmarks such as gold standard brain segmentations. Addressing this core limitation, we constructed the Baby Open Brains (BOBs) Repository, an open source resource comprising manually curated and expert-reviewed infant brain segmentations. Markers and expert reviewers manually segmented anatomical MRI data from 71 infant imaging visits across 51 participants, using both T1w and T2w images per visit. Anatomical images showed dramatic differences in myelination and intensities across the 1 to 9 month age range, emphasizing the need for densely sampled gold standard manual segmentations in these ages. The BOBs repository is publicly available through the Masonic Institute for the Developing Brain (MIDB) Open Data Initiative, which links S3 storage, Datalad for version control, and BrainBox for visualization. This repository represents an open-source paradigm, where new additions and changes can be added, enabling a community-driven resource that will improve over time and extend into new ages and protocols. These manual segmentations and the ongoing repository provide a benchmark for evaluating and improving pipelines dependent upon segmentations in the youngest populations. As such, this repository provides a vitally needed foundation for early-life large-scale studies such as HBCD.

## Main

Processing pipeline variability is a critical factor contributing to reproducibility challenges in neuroimaging research. When the same functional imaging dataset is analyzed by a variety of processing pipelines, different conclusions are drawn depending on which approaches were used ^1^. A variety of different processing stream decisions affect final conclusions, including pipeline components on both the structural and functional side ^2,3^. To support reproducible neuroimaging research, benchmarks must be identified for best standards and practices. One of these necessary benchmarks is gold standard manually defined brain tissue segmentations ^4^.

Nowhere are manually defined segmentations more needed than in studying the first 1000 days of life, a dynamically changing period of brain growth and development ^5,6^. Challenges inherent to the infant period due to this dynamic period of growth complicate accurate cortical and subcortical segmentation ^7^. The human brain undergoes considerable myelination through the first year of life, causing T1w scans (which enhance the signal of fatty tissue) and T2w scans (which enhance the signal of water) to show a contrast spin-inversion effect during this period. Existing studies remain limited due to protocols that varied considerably in processing mechanisms, including varied segmentation atlases.

Standardized infant segmentations atlases have become a critical need within research programs. The NIH has already invested $50+ million, and plans to invest hundreds of millions more, in the HEALthy Brain and Child Development (HBCD) study that promises to elucidate neurodevelopmental trajectories with unprecedented precision and rigor ^8^. This study promises to overcome sample size limitations highlighted by Marek et al. ^9^, filling a critical need for measuring true effect sizes for brain-wide associations relevant to early-life outcomes. Correct structural brain segmentations are essential to this promise, especially during the first 9 months due to the dynamic processes of growth and myelination occurring ^10–12^. Thus, an atlas is needed that supports the dynamic changes within this time period. Yet the availability of manually-corrected segmentations from anatomical MRI data across infancy is limited ^4^. Such corrections require considerable neuroanatomic expertise, expertise linking MRI landmarks to neuroanatomic borders, and are time-intensive, thus requiring considerable effort.

As field-wide momentum grows for reproducible research standards, a philosophy of open science is a necessary component of research best practices ^13^. Without transparent research, factors that contribute to low reproducibility rates cannot be examined. In this context, as underlying manual segmentations are an impactful part of processing pipelines, it stands to reason that these segmentations should themselves be open and transparent. The primary objective of this resource was to construct a set of manually curated and expert reviewed human infant brain segmentations that adhere to FAIR ^14^ data principles (Findable, Accessible, Interoperable, and Reusable). This repository can be used to assess existing pipelines and/or develop new ones, such as the recently presented BIBSNet algorithm that was trained on this repository ^15^. Early life segmentation algorithms already exist within the literature ^16–27^. However, many lack coverage across the whole-brain (eg. ID-Seg ^17^, MANTiS ^21^, iSEG challenges ^18^,

SDM U-net for subcortical ^16^, ANUBEX ^24^, SegSrgan ^23^), use only T1 or only T2 image inputs (eg. Infant Freesurfer ^19^, MCRIB-S^20^, ID-Seg^17^), or are specific to neonatal periods (VINNA^25^) and aren’t reliable across the full first years of life. As well, the underlying training data for those algorithms is often unavailable to the scientific community (iBEAT^26^). A lack of quality, publically available training data is a major limitation to improved infant segmentation pipelines, which is often pointed out by the developers of these algorithms themselves ^17,25^. Finally widespread disagreements among researchers can exist even for well-established areas like Wernicke’s area or the hippocampus ^28,29^. Therefore, making such manual corrections available via open repositories would subject such segmentations to broader exposure and review, improving the rigor and fidelity of the manual segmentations. Indeed, such work has already been performed extensively in adults and even in fetal tissue (ex. ^30–34^), and numerous segmentations have been made publicly available via repositories like OpenNeuro.org ^35^. Critically, this repository is open to new additions and changes, enabling a community driven resource that can improve over time and extend into new ages and protocol acquisitions.

## Results

### 71 BCP participants span 1-9 months of age

A schematic depicting the process of segmentation initialization, correction, and upload is shown in Figure 1. Markers performed image segmentation edits using ITK-SNAP ^36^ software. Markers attended trainings provided by the experts and had regular consultations with expert reviewers throughout the segmentation process. Initialized segmentations were overlaid on top of structural scans and manually edited. Markers utilized both the T1w and T2w scans to determine correct segmentation boundaries, such that there is one segmentation per session. Marker segmentations were reviewed by expert reviewers (EF/SS/JW/DA) and modified as needed. In total, segmentations were manually curated from 71 imaging visits across 51 participants. Of the 51 participants, 34 participants contributed one scan visit, 14 contributed 2 scans, and 3 contributed 3 scans to this set of segmentations. The age at scan ranged from 1-9 months old, with at least 6 scans at each month 1-8 (Figure 2). The demographics of the repository participants skewed White, non-Hispanic, and well-resourced (Figure 2), with 82% of the sample identifying as White, non-Hispanic and 96% of mothers having at least a college degree. Future versions and additions of the BOBs repository aim to make this distribution more diverse. Images selected for the repository represented best quality images based on visual review by the authors, which is considered a gold standard for quality assurance ^37^. The demographics of the 51 participants pulled for the repository did not differ statistically from the full BCP sample. Select neurodevelopmental measures, including the Mullen Scales of Early Learning, the Vineland Adaptive Behavior Scales, and subscales from the Infant Behavior Questionnaire - Revised, showed no differences between repository participants and the full BCP sample as well (Supplemental Table 1), suggesting that this participants in this repository can be considered representative of the larger BCP sample.

**Figure 1.**
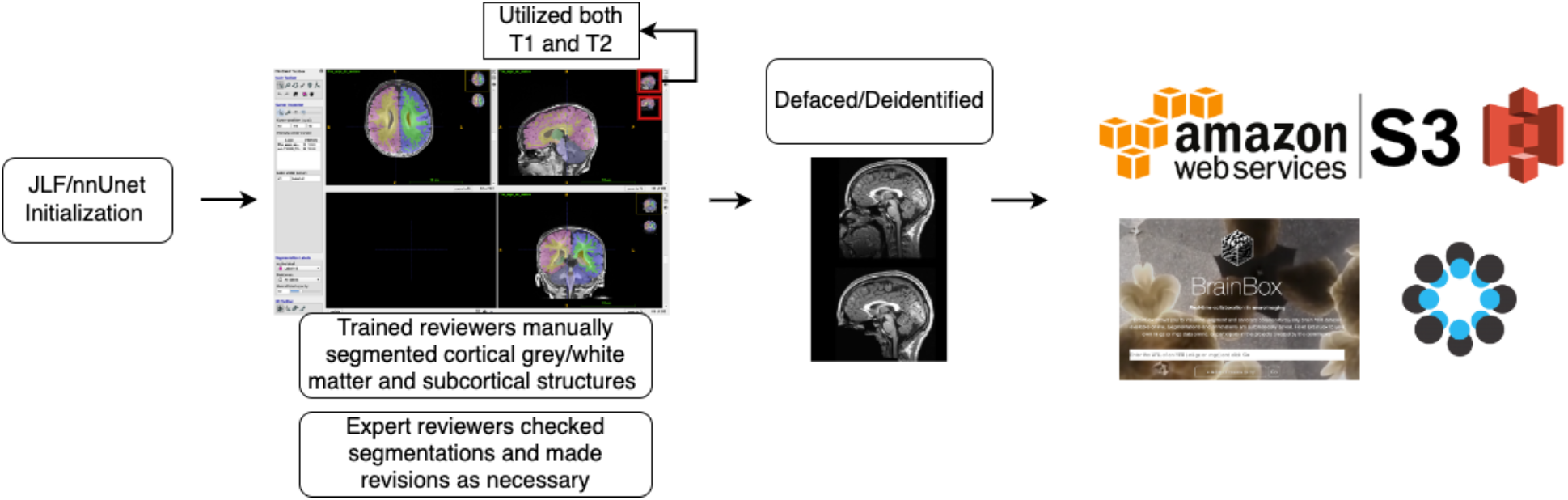
A schematic depicting the process of creating the repository. Segmentations were initialized with an automated processing pipeline and then manually corrected, utilizing both the T1 and T2 MRI images. Segmentations were then reviewed by expert reviewers who made revisions as necessary. These images were defaced and deidentified, uploaded to a public AWS S3 bucket, version controlled with DataLab, and then connected to BrainBox to allow researchers to review the repository. OSF acts as a hub to integrate the links to repository images, protocols, and any other future documentation created as the repository expands.

**Figure 2.**
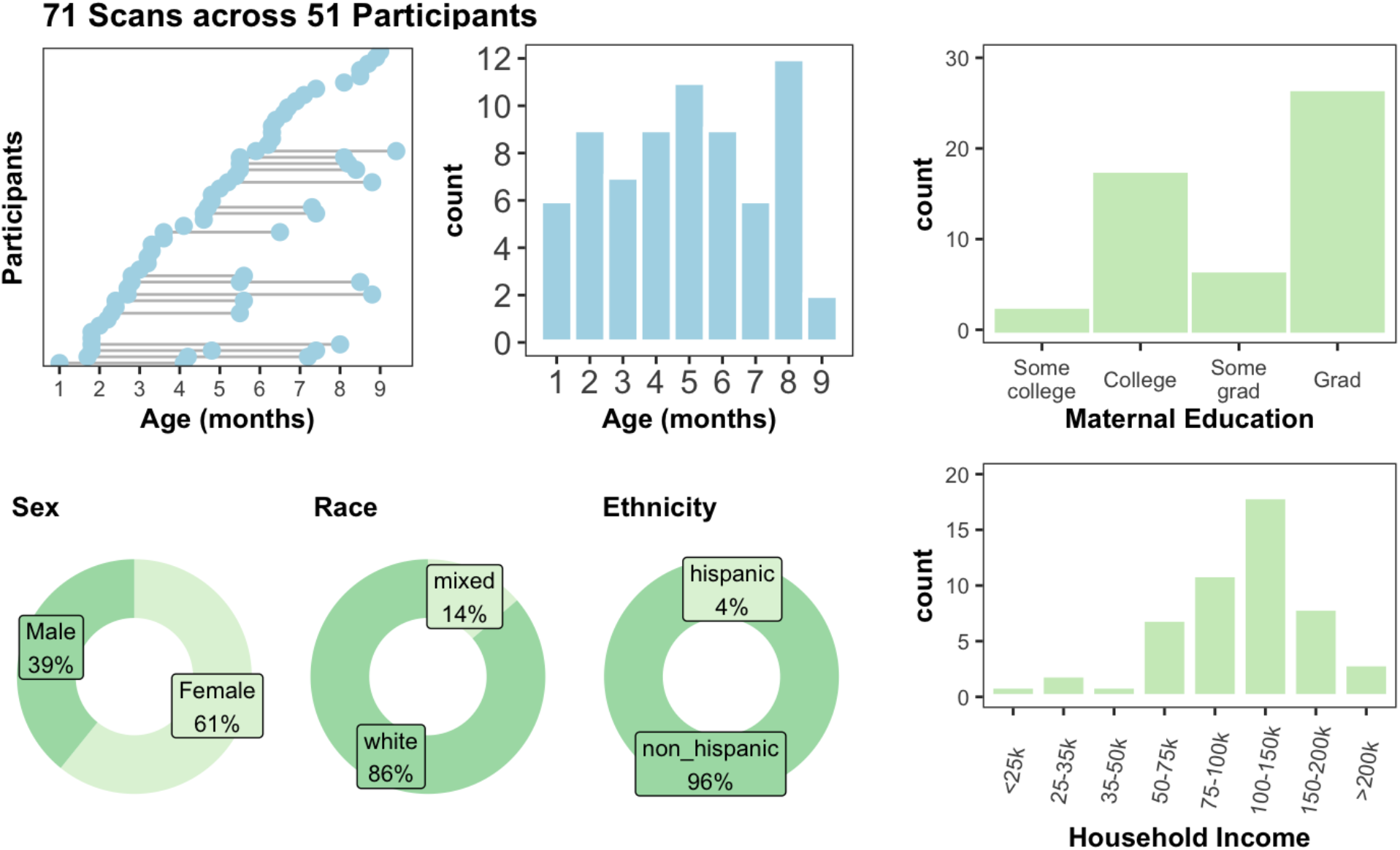
71 scans across 51 participants make up the segmentations in the repository. All come from the UMN site of the Baby Connectome Project (BCP), and span 1-9mo. The sample demographics skew white, non-hispanic, and well-resourced.

### The current BOBs repository comprises FreeSurfer-style segmentations for infants 1-9 months of age

These segmentations comprise cerebral gray/white matter and 23 subcortical structures. Uploaded segmentations went through several review stages before final approval, including at least one expert reviewer manually checking the segmentation. Leveraging both a T1w and T2w, care was taken to label white matter both affected and unaffected by the contrast spin-inversion effect. Diverging from FreeSurfer labels, the ventral thalamic boundary that separates thalamus from ventral diencephalon was defined by the hypothalamic sulcus ^38^. The hippocampal label was used to define the hippocampus proper, excluding the formation at the tail along the lingual gyrus, in order to be consistent with other infant literature ^39^. While we think evaluating whether the SOP is “right” or “wrong” may be beyond the scope of this paper, we chose such definitions in order to be more consistent with prior infant MRI literature ^39^. We welcome the community to inspect and refine existing segmentations to ensure that the “gold standard” benchmarks reflect a community gold standard.

Prior efforts towards developing automated segmentation pipelines lacked densely sampled, manually labeled training data that would be critical for early-life longitudinal studies like HBCD^4,8,10–12^. The Developing Human Connectome Project (dHCP) provides extensive anatomical segmentations that are largely restricted to neonatal and preterm infants ^22,40^. Such segmentations are derived from the T2w but do not use the T1w; while they can be used to develop automated segmentation pipelines ^16,23,24^, such pipelines may fail to generalize beyond the neonatal period. The iseg 2019 dataset comprises 39 datasets across multiple scanners and ages, and has produced 30 automated segmentation pipelines ^17,18^. However, manual segmentations are limited to gray matter, white matter, and CSF. The infant freesurfer dataset comprises data from a dozen infant sessions through the first two years of life ^41^, and helped develop infant freesurfer ^19^, but lacks the participant density of BOBs. The BOBs repository currently comprises the largest manually curated human infant brain segmentation dataset for the critical 1-9 month age range. This dataset proved vital in developing BIBSnet ^15^, an automated segmentation pipeline necessary for HBCD MRI data preprocessing, making the BOBs repository critical for infant pipeline development.

### Manual infant segmentation and curation requires tremendous effort

This age range in infancy and early childhood is a critical period of dynamic brain growth and development. 80% of brain growth occurs during the first 1000 days of life, including dramatic synaptogenesis, myelination, and other cellular processes ^42–44^. Aggregating over 100,000 participants from over 100 MRI studies, Bethlehem et al. found that brain development growth acceleration peaks at 7 months of age, with velocity highest around the first three years of life ^5^. Work by Alex et al confirmed this velocity peak and showed that these trajectories of growth are linked to cognitive and motor outcomes at 2 years of age and that these trajectories differ by sociodemographic factors and adverse birth outcomes ^6^.

As infant brains in this age range have increasing amounts of myelination in the white matter ^10^, referring to both T1w and T2w scans was critical to determining the extent of white matter. This brain growth is exemplified in a single infant in our repository across three ages in Figure 3. In this infant, there is visually dramatic development of image contrast within and across brain structures. This early time period shows rapid myelination, such that the older ages show much more myelinated white matter, especially along the major white matter tracts. Most dramatically at 5 months in this infant, there is an abundance of probably unmyelinated white matter that can be easily seen on the T2w image, but would be easily missed on the T1w image. Regardless of the cause, these developmental changes require considering both the T1w and the T2w images at this age group to fully capture the white matter and subcortical boundaries. This was especially critical in subcortical regions such as the basal ganglia, where boundaries might only be visible in either the T1w or the T2w, but not both. The blue arrows on each of the images exemplify regions that are better served by examining the T1w or the T2w but not both, such as the basal ganglia.

**Figure 3.**
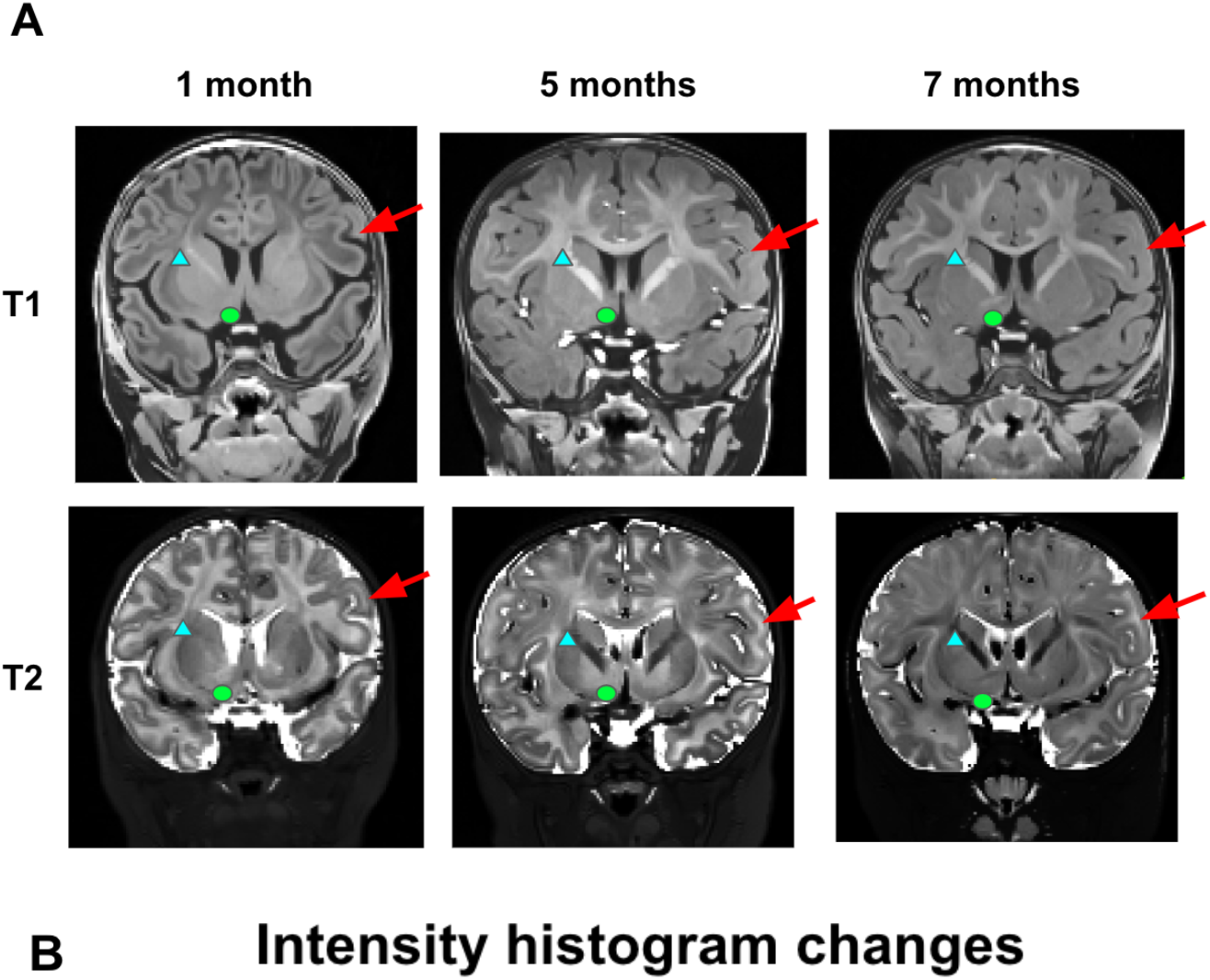
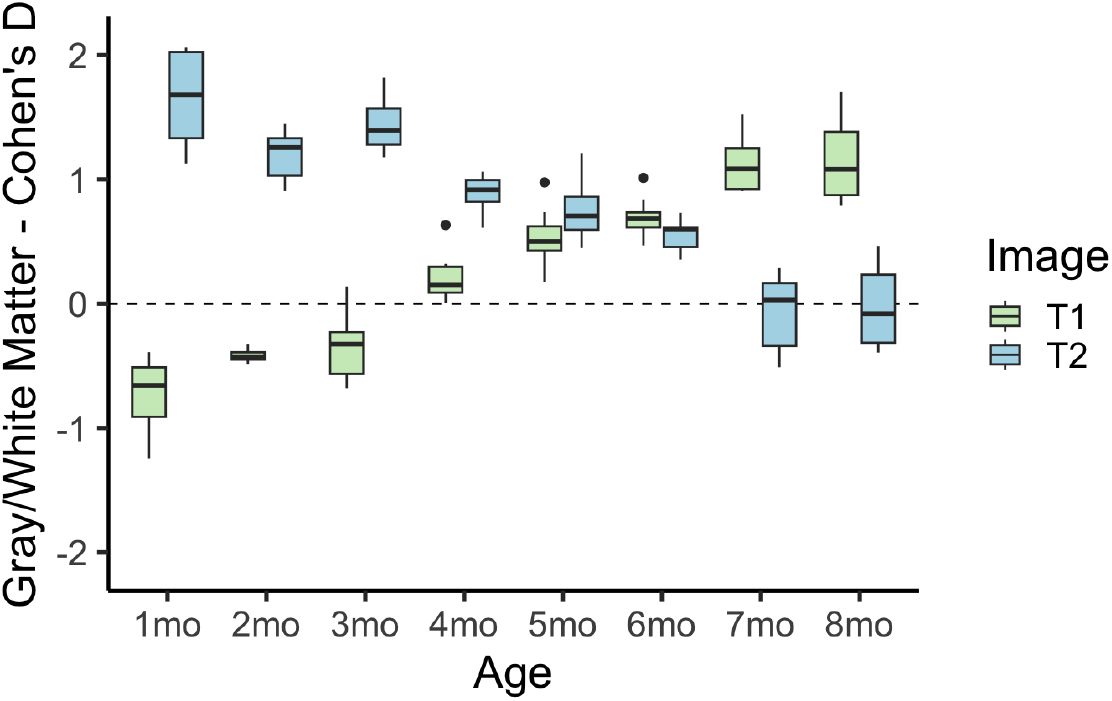
T1w and T2w images show dramatic developmental differences across the age range considered. (A) The selected images are from the same participant at three different ages, clearly depicting the transition from unmyelinated to myelinated white matter, and the differing image contrast intensities in the T1w vs. T2w at each age. Red arrows point out cortical gray/white matter changes, blue triangles point out internal capsule white matter changes, and green circles point out nucleus accumbens region changes (B) Cohen’s *d* values of white-gray matter differentiation are plotted for T1w and T2w MRI images. Considering both the T1w and the T2w images at this age group is critical to fully capture the white matter and subcortical boundaries.

### Manual segmentations show massive qualitative and quantitative improvement over initial Joint Label Fusion segmentations

Compared to initial Joint Label Fusion segmentations, created from the DCAN infant-ABCD-BIDS pipeline ^45^, manual segmentations show dramatic qualitative improvements. Initial segmentations had three major types of errors that were corrected by markers (see Figure 4). First, initial segmentations often created major errors in cortical folding patterns (Figure 4 top). The initial model may not account for differences between infant and adult image intensity, and this model failure may drive folding pattern segmentation errors. These errors required intensive edits to correct the basic gyri and sulci patterns. Additionally, due to the contrast spin inversion occurring at this age from myelination processes, labeling the full extent of unmyelinated white matter required extensive manual segmentation (Figure 4 middle). Automated segmentations often miss unmyelinated white matter, especially along the lateral surface of the brain where myelination processes occur later in development. Finally, as exemplified in Figure 3, subcortical regional intensities change dramatically over this time period, and thus subcortical regional boundaries often needed refining (Figure 4 bottom).

**Figure 4.**
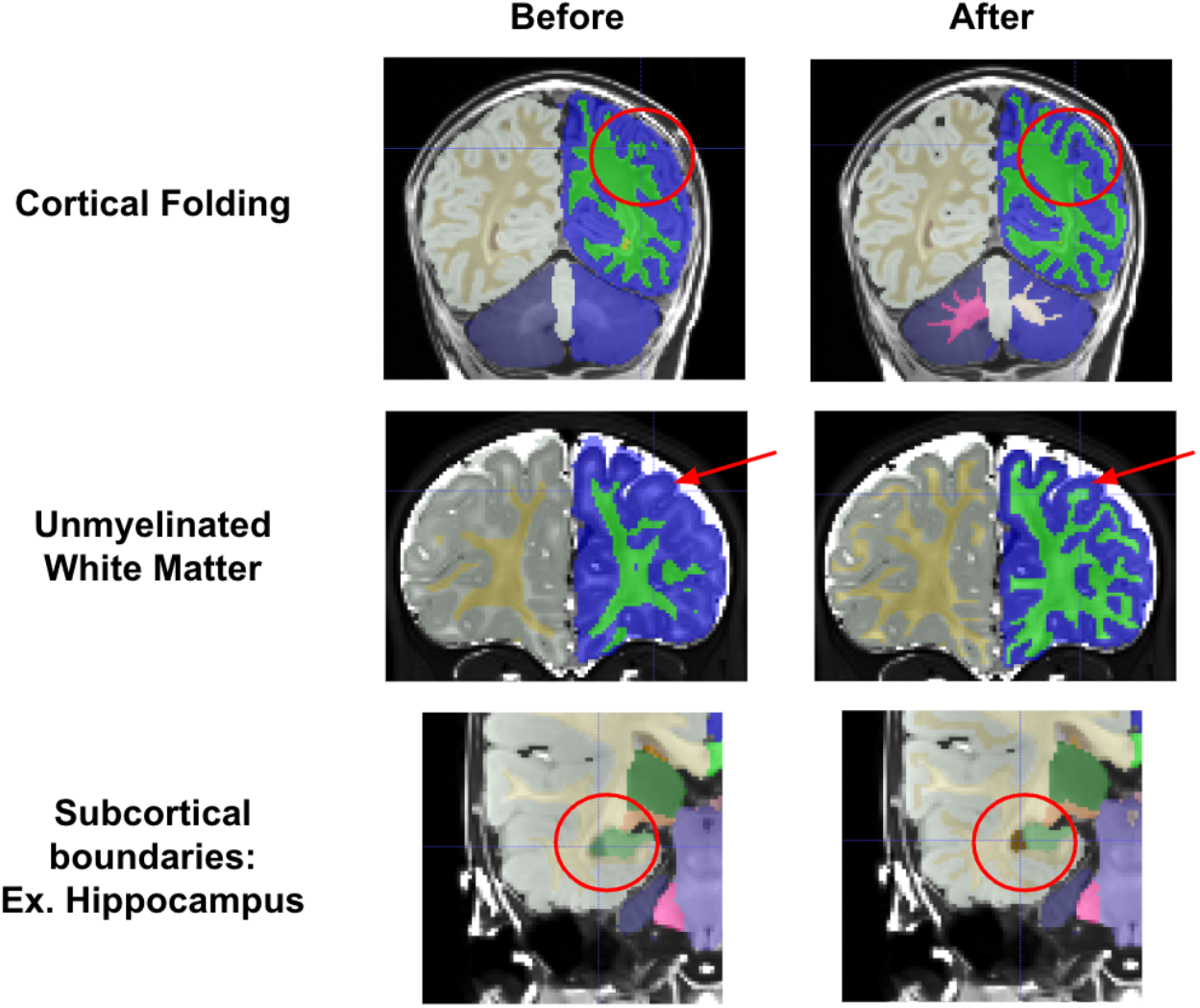
Manual segmentations show massive improvements over initial JLF segmentations. Three cases are demonstrated, showing that reviewers were able to correct errors such as cortical folding patterns, missing unmyelinated white matter, and incorrect subcortical boundaries.

### With S3 or BrainBox, researchers can add or refine segmentations

The BOBs repository is released and curated via the Masonic Institute for the Developing Brain Open Data Initiative, which leverages AWS S3 for storage and Datalad for version control. The structure of the repository provides all data in the BIDS standard, and with accessible links, which provides the ability for other applications (e.g., OpenNeuro) to leverage the repository as well without duplication. For example, leveraging this accessibility affords the ability for users to use BrainBox for visualization of the data ‘in place’, linked on our [ReadTheDocs](https://bobsrepository.readthedocs.io/en/latest/data_access/) page. To download repository segmentations to use or refine, researchers can access a download link on our [ReadTheDocs](https://bobsrepository.readthedocs.io/en/latest/data_access/) page, or use Cyberduck to have more control. The process to contribute refinements or new data to the repository is described below and outlined in detail on our [ReadTheDocs](https://bobsrepository.readthedocs.io/en/latest/data_access/). Briefly, contributors submit a form to the MIDB Open Data Initiative detailing their contributions. Contributions must include documentation on the segmentation edits or additions, and the images and segmentations should be submitted in BIDS format with Datalad version control. MIDB will then upload the documentation to the OSF page and the images and segmentations to the AWS S3 repository. All new contributions, including edits to existing segmentations will be tagged as new iterations of the repository, allowing researchers the freedom to select any dataset’s iteration.

## Discussion

The BOBs repository addresses the need for openly available manually corrected segmentations of MRI data during the earliest periods of life ^4^. Such a resource is critical for developers wishing to create processes for accurate automated segmentations. The curation of such a repository requires considerable neuroanatomic expertise, including knowledge of anatomical MRI landmarks for accurate segmentations. Until now, the labor and considerable effort required to conduct such work has left much of the methods development without a ‘gold standard’ or benchmark dataset. This lack of proper benchmarks has limited the ability for pipeline developers to generalize infant processing pipelines and ensure the effectiveness of different pipelines across infant age groups, which has subsequently led to constrained pipelines tuned for particular ages.

BOBs manual segmentations will provide a benchmark for evaluating and improving automated segmentations. As infant neuroimaging expands, the research community will observe an exponential increase in MRI segmentation approaches. Already at least a half dozen early life segmentation algorithms exist within the literature ^16–24^; however, few have tested segmentations across the early life age span and that cover the whole-brain and incorporate a wide-breadth of labels beyond gray matter, white matter, and CSF. Combining expert and community review, the BOBs repository provides a unique foundational benchmark for evaluating and improving image segmentation methods, as well as expanding their scope towards more comprehensive segmentations. Such benchmarks standardize methods development as methods researchers can evaluate segmentation performance and validate tool capability. Such a benchmark standard for performance evaluation facilitates best practices and standards in infant neuroimaging.

Data repositories operating under findable, accessible, interoperable, and reproducible (FAIR) principles enable best standards and practices for infant MRI pipeline development. The BOBs repository provides a centralized location for housing manually segmented infant MRI data that adheres to FAIR principles, and informs researchers on the providence and fidelity of the segmentations. BrainBox ^46^ and version control with Datalad ^47^ enable community users to review BOBs repository datasets to ensure that datasets meet robust community-wide standards, creating a robust gold standard for future MRI pipeline development. Through MIDB’s Open Data Initiative, researchers can review and submit edits to segmentations, as well as make informed decisions regarding selecting manual segmentations knowing who created them and why. Revised segmentations and new datasets will be incorporated as new iterations according to the process outlined in the Results section. The initial dataset for the BOBs repository includes infant segmentations from 1 to 9 months that have been curated by a team comprising four experts. Making this repository FAIR ensures that infant neuroimaging pipeline development will fulfill calls for reproducible neuroscience research ^48^.

Future directions include expanding FreeSurfer-style segmentations and adding new segmentations to BOBs. While these 71 scans represent the largest known repository for manually curated whole-brain FreeSurfer-style segmentations aligned with T1w and T2w data, we anticipate the need for even more ground truth segmentations in the future. These infants were all collected using a single protocol on a single scanner. Variations in acquisition protocol, scanner or platform may limit the application of algorithms across the community and big data consortia like HBCD. In addition, infant studies often examine age ranges beyond 9 months.

Therefore, future efforts may add to the repository collection in order to provide benchmarks that generalize across research sites and the early human lifespan. In addition, the infant brain imaging community remains interested in neuroanatomical regions beyond FreeSurfer-style segmentations. Alternative approaches for surface generation like MCRIBS and DHCP may rely on different brain segmentations, requiring a new set of labels to be incorporated in pipelines.

Thalamic, amygdala, and hippocampal subregion development remains of great interest in the infant community. Such segmentations require higher resolution brain scans and curated manual segmentations to benchmark algorithms. Future efforts will expand the repository with additional curated data that provide generalizable benchmarks across different brain region definitions.

These algorithms will form a necessary foundation for early-life large-scale studies such as HBCD. Automated MR processing pipelines specifically designed for early development are necessary to allow large-scale studies such as HBCD to create MR outputs unconfounded by age. With the BOBs resource providing a foundational benchmark to evaluate and improve these processing pipelines, HBCD and other future early-life neuroimaging studies will be well-equipped to provide the promised knowledge of nuanced neurodevelopmental trajectories and their complex environmental interactions.

## Methods

### The repository is comprised of Baby Connectome Project (BCP) anatomic and segmentation MRI data

The Baby Connectome Project (BCP) is a longitudinal neuroimaging study in infants 0-5 years old. Detailed methodology has been described previously (Howell et al. 2019), but briefly, infants were recruited from departmental research participant registries based on both state-wide birth records and the broader communities. Infants were eligible for the BCP if they 1) were born at a gestational age of 37–42 weeks, 2) had a birth weight appropriate for gestational age, and 3) had an absence of major pregnancy and delivery complications. For this repository, 71 MRI visits with good quality data from infants 1-9 months old scanned at the University of Minnesota were used. MRI data was collected using a 32-channel head coil on a Siemens 3T Prisma scanner and included high resolution T1w (MPRAGE: TR 2400ms, TE 2.24ms, TI

1600ms, Flip angle 8°, resolution = 0.8 × 0.8 × 0.8 mm3) and T2w (turbo spin-echo sequences: turbo factor 314, Echo train length 1166ms, TR 3200ms, TE 564ms, resolution = 0.8 × 0.8 × 0.8 mm3, with a variable flip angle) structural scans collected during natural sleep.

### Segmentations were initialized using two different segmentation pipelines

As a starting point for manual reviewers, segmentations were run through one of two segmentation pipelines. Some segmentations were initialized from a joint label fusion (JLF) pipeline ^49^, and then manually curated. However, such a procedure required many hours of manual curation as these initializations required much coarser edits. Therefore, these initial manual segmentations were used to train “BIBSNet” ^15^, a deep neural network built using nnU-Net ^50^ and SynthSeg ^27^. Using BIBSNet, other segmentations were initialized and then manually curated. Iteratively using BIBSNet prototypes as a starting point saved many hours of work, as the prototypes were much more accurate starting points than the JLF pipeline. In both pipelines, Advanced Normalization Tools (ANTs) was used to perform denoising and N4 bias field correction and T1w and T2w images underwent a rigid-body realignment to remove distortions and improve image quality for the reviewers.

### Markers curated segmentations according to a standard operating protocol

Markers performed image segmentation edits using ITK-SNAP ^36^ software. Initialized segmentations were overlaid on top of structural scans and manually edited. Markers utilized both the T1w and T2w scans to determine correct segmentation boundaries. As infant brains in this age range have increasing amounts of myelination in the white matter, referring to both T1w and T2w scans was critical to determining the extent of white matter. For each brain, the cortical surface and the gray-white matter boundary were edited first and reviewed. Subcortical regions were then edited, including the lateral ventricles, inferior lateral ventricles, cerebellum white matter, cerebellum cortex, thalamus, caudate, putamen, pallidum, amygdala, hippocampus, nucleus accumbens, third ventricle, fourth ventricle, and brainstem. Segmentations were done in phases, with the lateral ventricles, third ventricle, and fourth ventricle segmented first, the nucleus accumbens, caudate, putamen, and pallidum second, the brainstem, thalamus, and cerebellum third, and then the amygdala and inferior lateral ventricles last. The hippocampus was segmented separately, either before or after the rest of the subcortical segmentations.

Definitions for the boundaries of these regions were pulled from previously published definitions ^51–53^. A full SOP of subcortical boundaries was created and can be found on the OSF site (https://osf.io/wdr78/) as well as the ReadTheDocs page (https://bobsrepository.readthedocs.io).

### Approved anatomic MRI data were deidentified, defaced, and uploaded to a public S3 bucket

Final data was stripped of identifying information and formatted into BIDS format. To deface images, T1w and T2w images were run through PyDeface using MNI infant templates as well as a custom infant mask (https://cdnis-brain.readthedocs.io/defacing/), which masked out facial features from the scans. Final deidentified and defaced images and segmentations were uploaded to a public AWS S3 bucket and version controlled with DataLad. This bucket was then linked to BrainBox (https://brainbox.pasteur.fr/), which allows users to review the repository online.

**Supplemental Table 1.** No differences were seen on selected neurodevelopmental scores between the participants selected for the BOBs repository and the full BCP sample from which they were selected.

|  | BOBs | BCP | p-value |
| --- | --- | --- | --- |
| Mullen Early Learning Composite Score (SD) | 103.09 (12.67) | 104.98 (14.69) | 0.34 |
| Vineland Adaptive Behavior Composite (SD) | 96.96 (7.66) | 99.26 (9.26) | 0.07 |
| Infant Behavior Questionnaire - Revised: Smiling and Laughing (SD) | 4.61 (0.98) | 4.59 (1.10) | 0.93 |
| Infant Behavior Questionnaire - Revised: Fear (SD) | 2.35 (0.82) | 2.19 (0.82) | 0.29 |
| Infant Behavior Questionnaire - Revised: Duration of Orienting (SD) | 3.64 (1.12) | 3.65 (1.11) | 0.96 |

<sup>^</sup>For the Vineland and IBQ-R, as differences are seen in scores across ages, BOBs participants were compared to only those in the BOBs age range. For the Mullen, the age standardized composite score was used, so BOBs participants were compared to all BCP participants. There were also no differences when just compared between those in the BOBs age range.

## Supporting information

Supplemental Segmentation SOP

## Acknowledgements

The authors would like to thank additional markers and coordinators who contributed to this repository, including Henrique A. Caldas, BettyAnn Chodkowski, Katie Day, Ekomobong Eyoh, Brayton Hall, Alexandra Harper, and Sarah Kuplic. This work was supported by Bill & Melinda Gates Foundation INV-015711 (M.D.R., J.T.E., C.D.S., D.A.F.) The UNC/UMN Baby Connectome Project was supported by NIMH R01 MH104324 and NIMH U01 MH110274. EAK is supported by a National Research Service Award (NRSA) T32 training grant (T32-NS109604). SMS and TKMD are supported by the National Science Foundation Graduate Research Fellowship Program (SMS: 2237827; TKMD: 2020295366). DAF is supported by NIDA U01DA041148, NIDA U24DA055330, NIMH R01MH096773, NIMH R01MH125829, and NIMH R37MH125829.

## Notes

### Competing Interest Statement

D.A. Fair has ownership Interest in FIRMM software, Turing Medical Inc; M. Bagonis, K. Barrett, B. Bower, D. Goradia, L. Heisler-Roman, C. Lucena, M. Myricks, and P. Narnur all worked for PrimeNeuro at the time of their contributions to this manuscript.

### Summary of Updates

Competing Interest Statement was updated to remove an author who was mistakenly added.

