## Supplemental Segmentation SOP for "Baby Open Brains: An Open-Source Repository of Infant Brain Segmentations"

### Infant-specific notes on the Standard Operating Procedures (SOP) for manual segmentation

#### Subcortical Segmentation Workflow

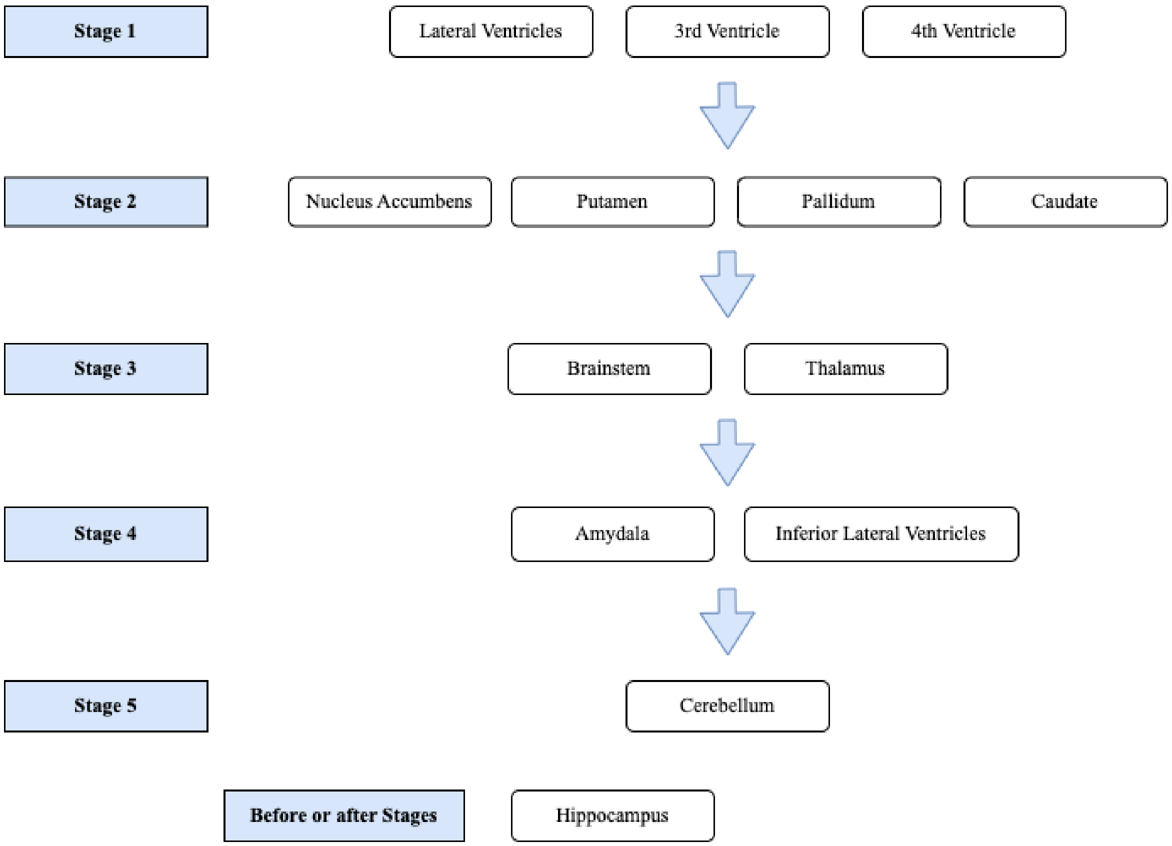

### Common Abbreviations

Cerebellum = Cb
Putamen = Put
Nucleus Accumbens = NA
Globus pallidus = GP (GPe = external, i=internal)
Hippocampus = HC
Amygdala = Amy
Caudate = Caud
Optic Chiasm = OC

### General Principles

The majority of the segmentation definitions here are based on adult segmentation protocols, with adjustments and refinements for infants. Thus, this protocol refers to those adult segmentation protocols for general information about the structures. This protocol is not intended to be a stand-alone document for those who have never segmented brain tissue before, it is intended to convey the nuances of infant brain segmentation and the nuances of this protocol specifically. As such, you will need to refer to these documents as well for general overviews:

Rushmore, R. J., Sunderland, K., Carrington, H., Chen, J., Halle, M., Lasso, A., ... & Makris, N. (2022). Anatomically curated segmentation of human subcortical structures in high resolution magnetic resonance imaging: An open science approach. Frontiers in Neuroanatomy, 16, 894606.

- Most of the structures refer to this citation
- The supplemental material includes a manual that was often utilized by our markers
    - Manual for Segmentation of Subcortical Structures (Data Sheet 1) https://www.frontiersin.org/articles/10.3389/fnana.2022.894606/full#supplementary-material

Wedig, M. M., Rauch, S. L., Albert, M. S., & Wright, C. I. (2005). Differential amygdala habituation to neutral faces in young and elderly adults. Neuroscience Letters, 385(2), 114-119.

- For amygdala

Watson, C., Andermann, F., Gloor, P. M. D. P., Jones-Gotman, M., Peters, T., Evans, A., ... & Leroux, G. (1992). Anatomic basis of amygdaloid and hippocampal volume measurement by magnetic resonance imaging. Neurology, 42(9), 1743-1743.

Jack Jr, C. R. (1994). MRI-based hippocampal volume measurements in epilepsy. Epilepsia, 35, S21-S29.

- For hippocampus

Label Numbers (Derived from Freesurfer)

| **Structure** | **Label** |
|---|---|
| Left White Matter | 2 |
| Left Gray Matter | 3 |
| Left Lateral Ventricle | 4 |
| Left Inferior Lateral Ventricle | 5 |
| Left Cerebellum White Matter | 7 |
| Left Cerebellum Cortex | 8 |
| Left Thalamus | 10 |
| Left Caudate | 11 |
| Left Putamen | 12 |
| Left Pallidum | 13 |
| 3rd Ventricle | 14 |
| 4th Ventricle | 15 |
| Brainstem | 16 |
| Left Hippocampus | 17 |
| Left Amygdala | 18 |
| Left Nucleus Accumbens | 26 |
| Left Choroid Plexus | 31 |
| Right White Matter | 41 |
| Right Gray Matter | 42 |
| Right Lateral Ventricle | 43 |
| Right Inferior Lateral Ventricle | 44 |
| Right Cerebellum White Matter | 46 |
| Right Cerebellum Cortex | 47 |
| Right Thalamus | 49 |
| Right Caudate | 50 |
| Right Putamen | 51 |
| Right Pallidum | 52 |
| Right Hippocampus | 53 |
| Right Amygdala | 54 |
| Right Nucleus Accumbens | 58 |
| Right Choroid Plexus | 63 |

A couple general points about this segmentation workflow:

- In this protocol, we follow neuroanatomical standards, such that the left side of an image is the right hemisphere, and the right side of an image is the left hemisphere. This is the standard in the ITK-SNAP software used in the manual segmentations as well.
- Myelinated and unmyelinated white matter were not differentiated for this protocol, and were both segmented as white matter
- Limitation: The ventral diencephalon (including substantia nigra) is not explicitly segmented in this protocol, so it is only implicitly defined by the boundaries of surrounding structures that were segmented in this protocol. Similarly, vermis (see Cerebellum section) and WM-hypointensities were Freesurfer labels that were not changed if they existed, so are inconsistently used across participants.

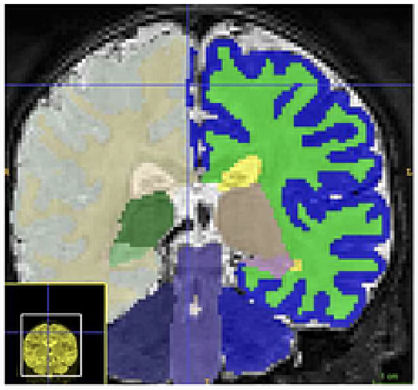

- Keep an eye out for swapped left/right hemisphere labels around the midline as this is a common error we observe from segmentations produced by our deep learning models. This can impact both the gray/white matter and subcortical structures. In the image to the right, the gray matter of the right hemisphere (gray, displayed on the left in this image) is inappropriately labeled as left hemisphere gray matter (blue) around the midline
- ROI delineations can look “choppy” if they are only corrected for in one plane of view, so make sure to check all fields of view to reduce this. In the example below, the coronal plane was nicely corrected, but the sagittal and axial views look “choppy.”

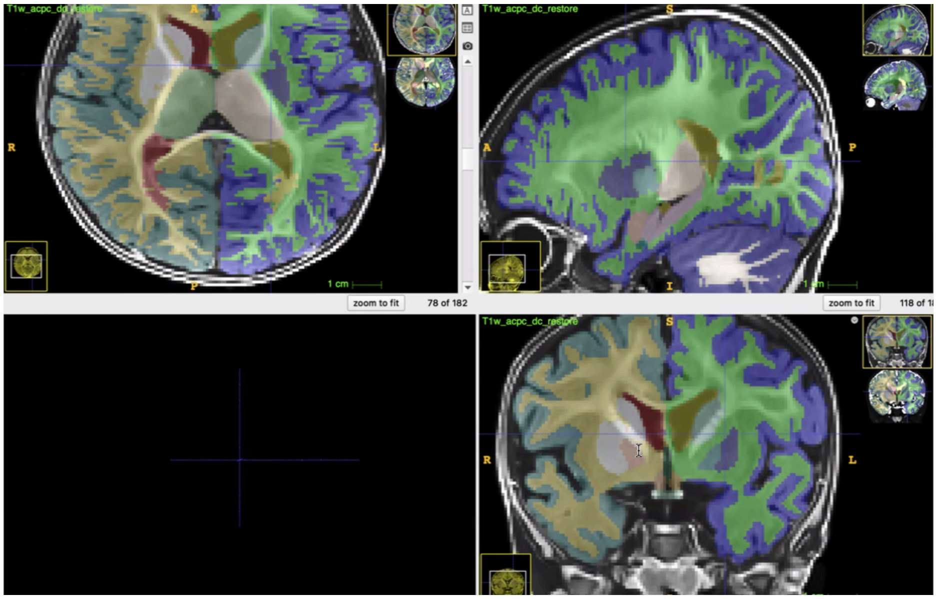

- If a label is missing entirely (eg the label for the choroid plexus has been noted as entirely missing in some segmentations), then you can add the label in ITKSnap in order to label that region

### Region-specific SOPs

#### Gray (GM) and White Matter (WM)

Use both the T1 and T2 images to make sure that gray matter and white matter are accurately labeled across all ages and tissue types.

Some Common Issues:

- Cortical folding patterns
  - Gyri are missing and not labeled as brain matter
  - White matter is under or over segmented such that the folding pattern is grossly mislabeled (see example to the right)

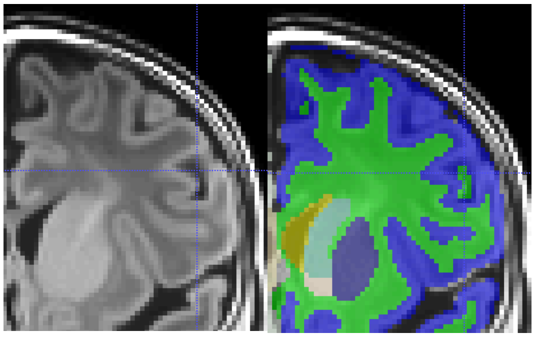

- Unmyelinated white matter missing
  - Much of the white matter that is missing is because the automated segmentation algorithms miss unmyelinated white matter (see example below).
  - Especially in the younger ages, be sure to use the T2 to capture the entirety of the unmyelinated white matter. In ages 4 - 6 months or so, both the T1 and T2 will be very necessary as different regions in the brain will comprise both myelinated and unmyelinated white matter, which may not be easy to see on a single image modality.

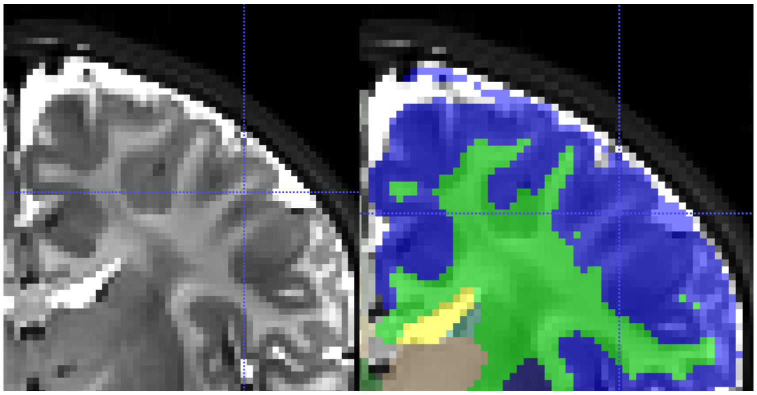

- Pay special attention to white matter correction in occipital cortex and motor cortex (pre/post-central gyri) as these areas typically require the most

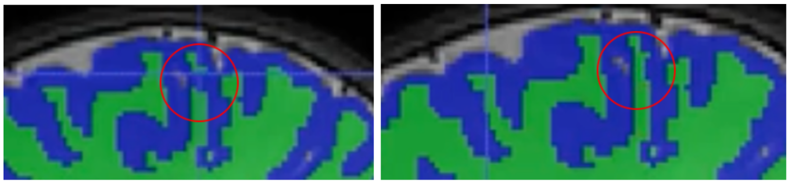

attention/edits. The screenshots to the right in the motor cortex are one example that would cause issues for later cortical surface reconstruction.

- Make sure to segment white matter of internal capsule - this was often missed

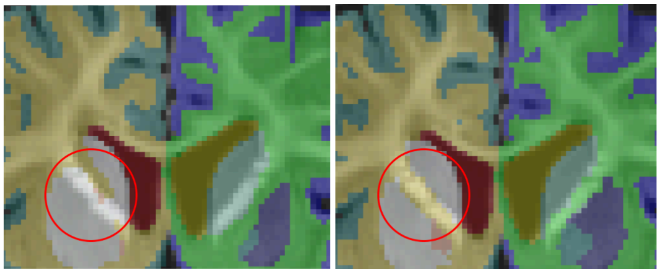

### Left/Right Lateral Ventricle

Based on “The lateral ventricle” section in (Rushmore et al. 2022) - see the supplement for more detailed figures.

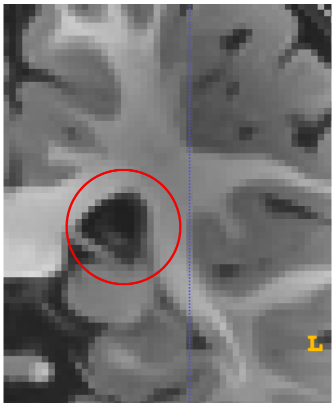


- Boundaries are relatively clear as the ventricle is relatively distinct from adjacent white matter and regions. Take care that the boundary with the third ventricle is clear (see Third Ventricle section in this SOP). When the hippocampus appears (when the fornix is disconnected) then the ventricle splits from the lateral ventricle into inferior lateral ventricle. See figure to the right for the slice where that happens. The red circle indicates the lateral ventricle.
- Within the lateral ventricle are “wisps” of gray matter called choroid plexus - these have their own label as well that should be corrected. A small amount can be seen in the figure to the right. A substantial amount of choroid plexus is seen in the youngest infants, but be careful to distinguish choroid plexus from surrounding structures (ex. white matter, caudate). The choroid plexus may not be visible at all brightness levels, so be sure to adjust the contrast such that the choroid plexus is visible while segmenting.

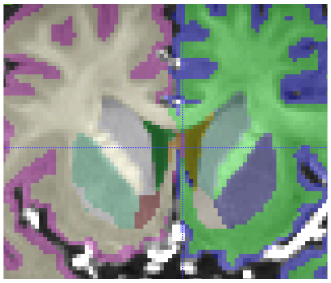

- A few common issues can occur in the lateral ventricle:
  - There may be instances where the choroid plexus label is missing entirely. When this happens, you can simply add a new label in ITK-snap in order to segment it.
  - Another common issue is that the choroid plexus is over-segmented at the midline (see figure to the right)

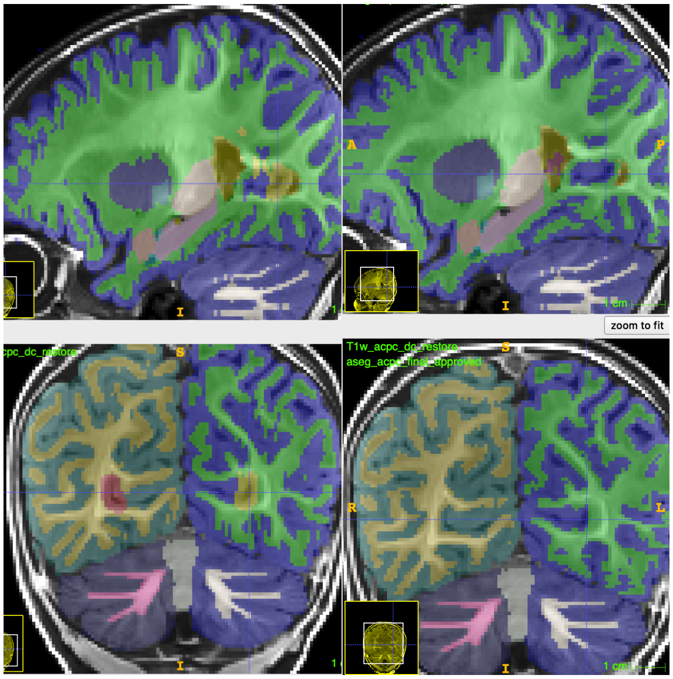


  - There are cases where the lateral ventricle is over-segmented posteriorly. In the example to the right, the crosshairs show an area that is labeled as ventricle, but is actually gray matter. This is then corrected in the second screenshot:

Lateral Ventricle Figures

| Age | T1 | T2 | Segmentation |
|---|---|---|---|
| 1mo | 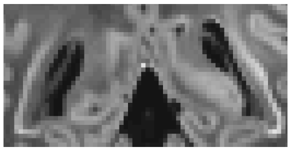 | 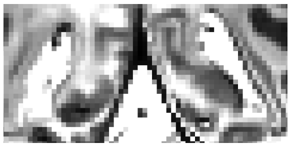 | 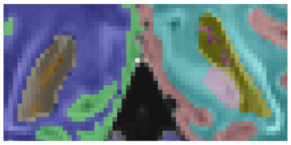 |
| 4mo | 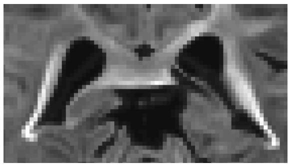 | 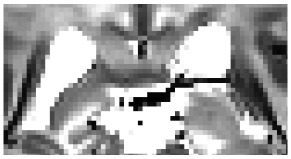 | 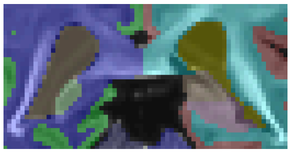 |
| 6mo | 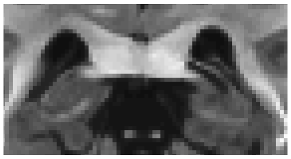 | 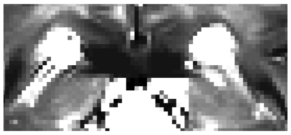 | 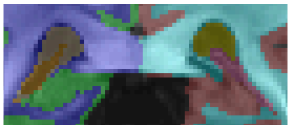 |
| 8mo | 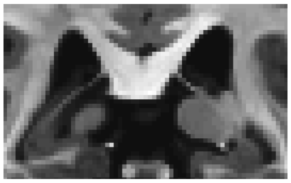 | 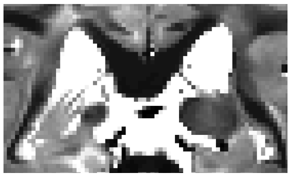 | 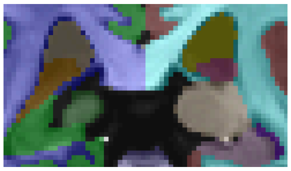 |

Choroid Plexus Figures

| Age | T1 | T2 | Segmentation |
|---|---|---|---|
| 1mo | 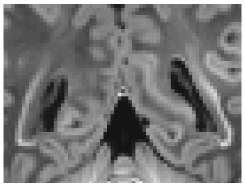 | 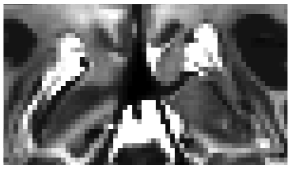 | 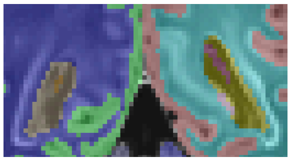 |
| 4mo | 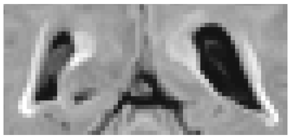 | 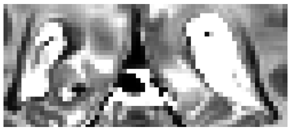 | 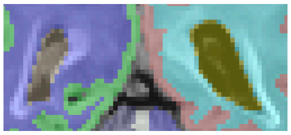 |
| 6mo | 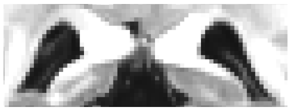 | 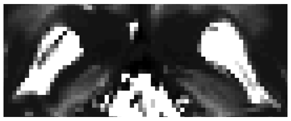 | 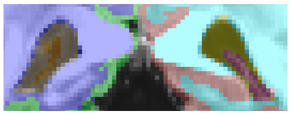 |

| | | | |
|---|---|---|---|
| 8mo | 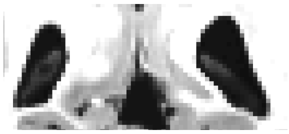 | 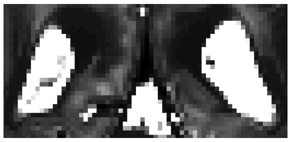 | 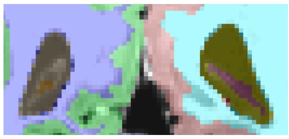 |

Choroid Plexus Contrast Figures

| | 1mo Bright | 1mo | 1mo Dark |
|---|---|---|---|
| T1 | 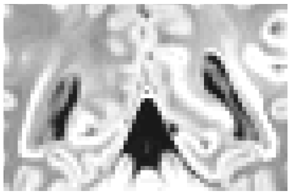 | 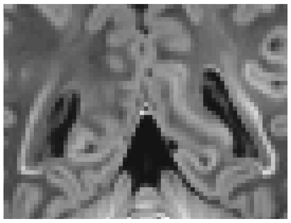 | 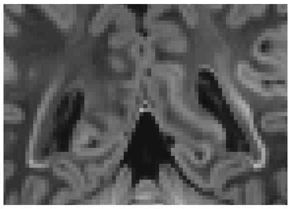 |

### Third Ventricle

Based on “The third ventricle” section in (Rushmore et al. 2022)

- The anterior border is defined by the small indent in optic chiasm as seen in figure below (L, no indent, R, next slice with indent, as indicated by the gridlines). This quickly turns into a “teardrop” shape. A common issue seen for this structure is that the 3rd ventricle doesn’t start earlier enough at this “teardrop shape”

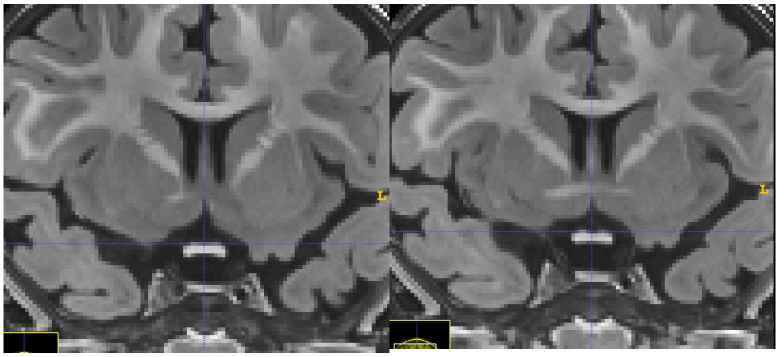

- On the figure to the right, where the ventricle starts to open up, include up to the gray matter structure at the top of the ventricle (including a little bit of the choroid plexus below it). Pictured here on T1, but the T2 (not pictured) will start to be helpful as well. On T2 most of it will be dark, with only the little bit of choroid plexus that will be included as light.

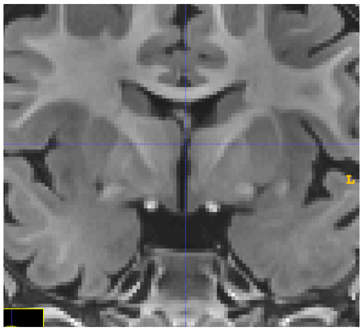

- The T2 is helpful in determining the posterior end of the third ventricle. Include only the dark region (see the grid line on the figure to the right) until it disappears.

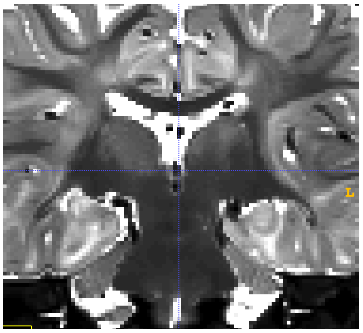

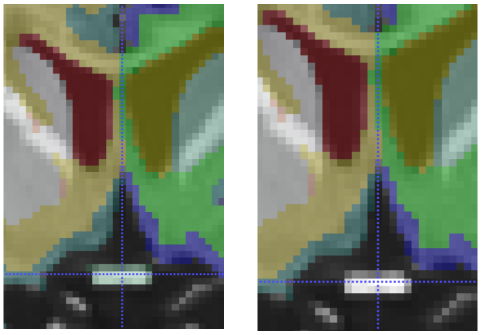

- A common issue in the 3rd ventricle is over-segmentation onto the optic nerve (as seen on left, correct on right)

Third Ventricle Figures

| Age | T1 | T2 | Segmentation |
|---|---|---|---|
| 1mo | 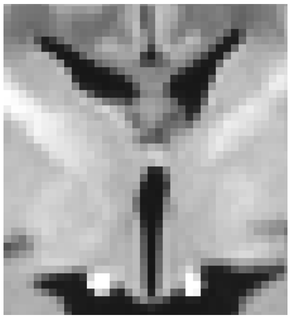 | 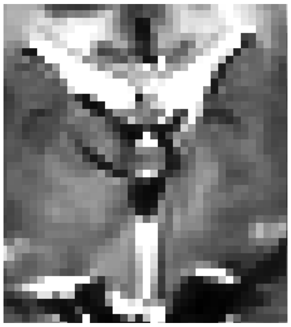 | 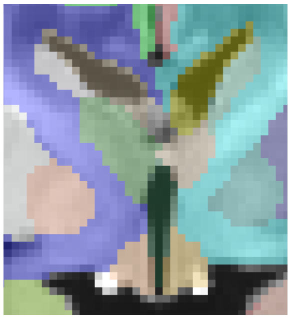 |
| 4mo | 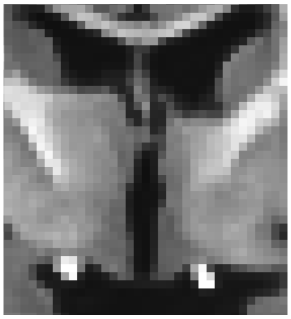 | 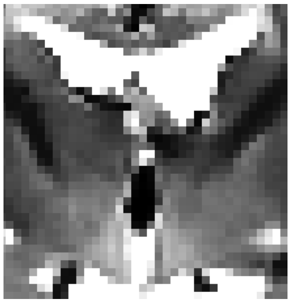 | 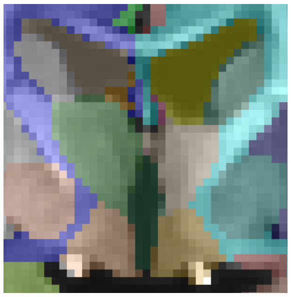 |
| 6mo | 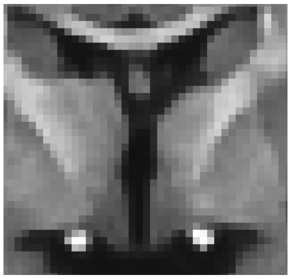 | 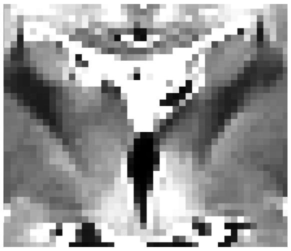 | 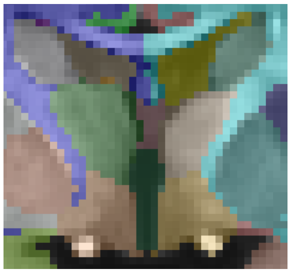 |
| 8mo | 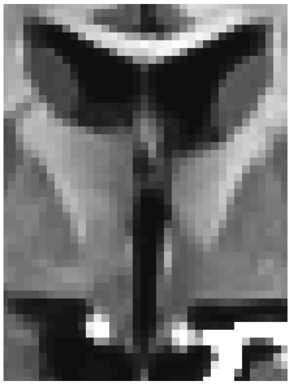 | 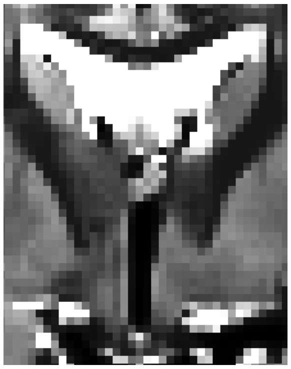 | 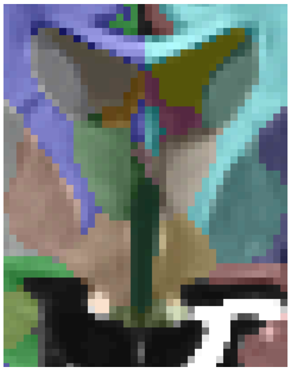 |

Teardrop shape Figures

| Age | T1 | T2 | Segmentation |
|---|---|---|---|
| 1mo | 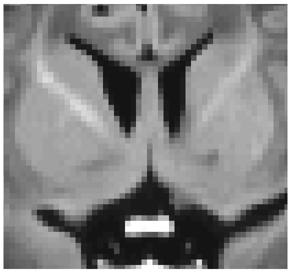 | 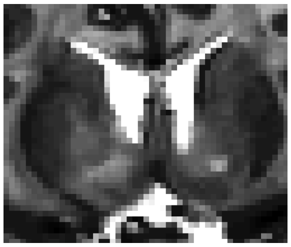 | 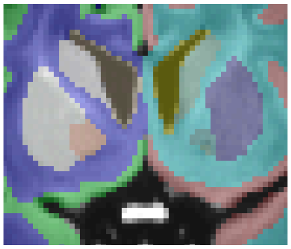 |
| 4mo | 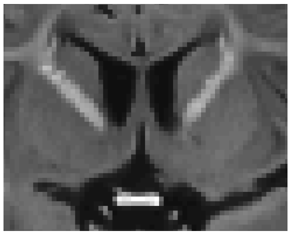 | 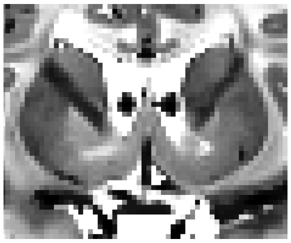 | 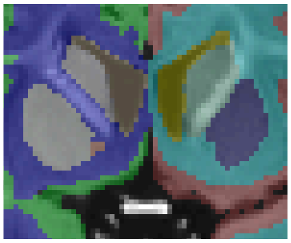 |
| 6mo | 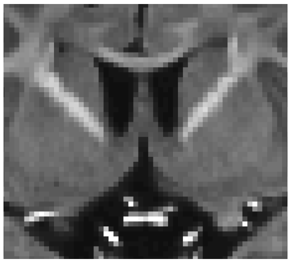 | 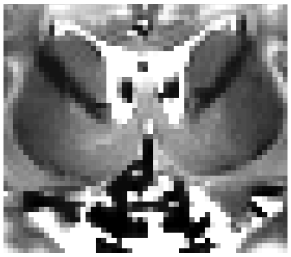 | 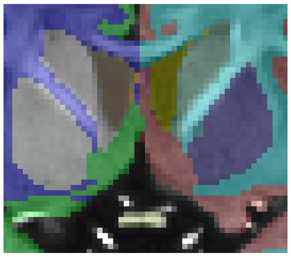 |
| 8mo | 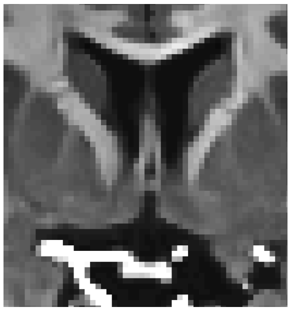 | 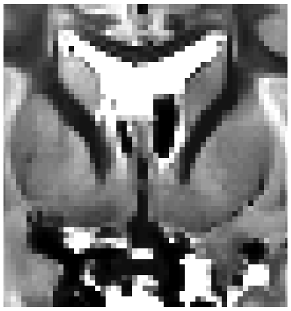 | 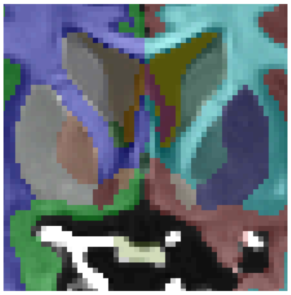 |

### Fourth Ventricle

Based on “The fourth ventricle” section in (Rushmore et al. 2022)

- Fourth ventricle starts at the dot in the brainstem that is used in the brainstem definition (“cerebral aqueduct of Sylvius”, see grid line on figure to the right). It continues down until the “obex” (red circle) where the brainstem ends.
- As it continues, it splits in two (top and bottom in coronal). Be sure to include both of these sections. See grid line below in the inferior section. Eventually the inferior portion disappears and the superior portion splits into left and right in coronal as well.

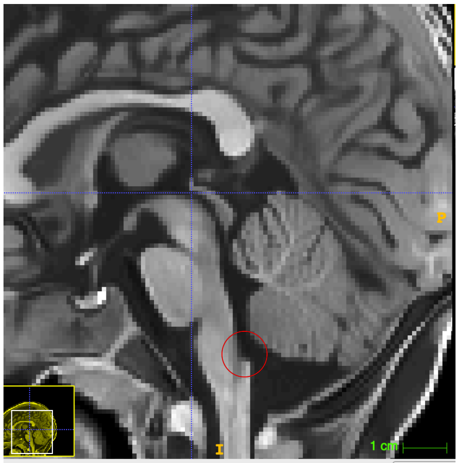


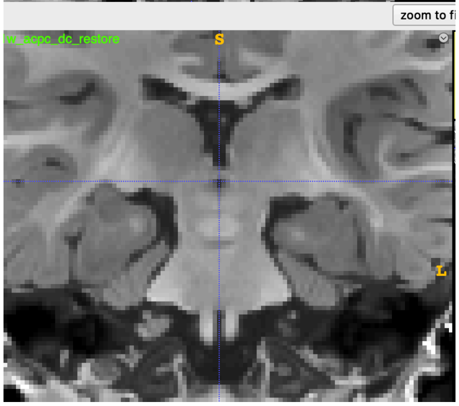


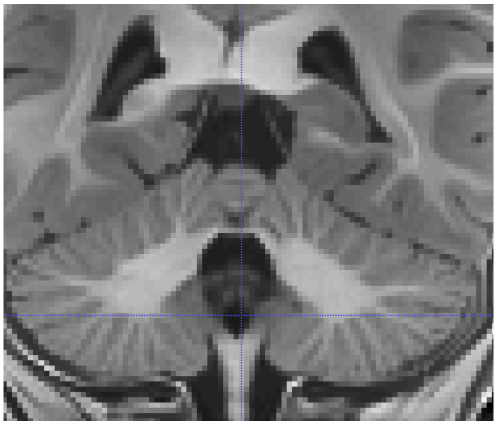

#### Fourth Ventricle Figures

| Age | T1 | T2 | Segmentation |
|---|---|---|---|
| 1mo | 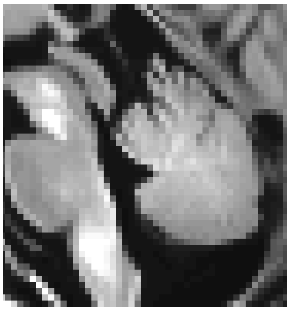 | 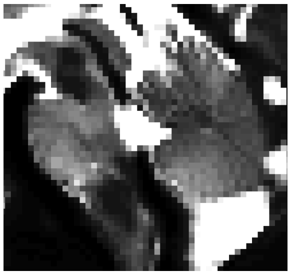 | 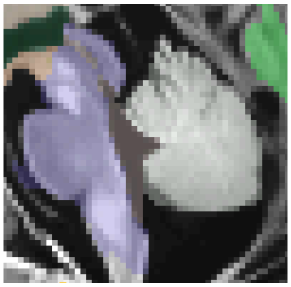 |

| | | | |
|---|---|---|---|
| 4mo | 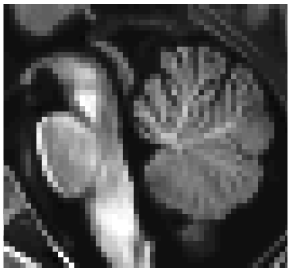 | 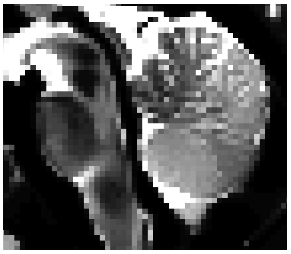 | 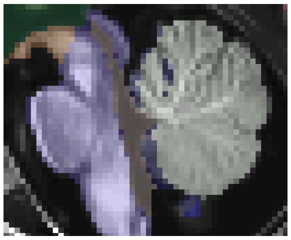 |
| 6mo | 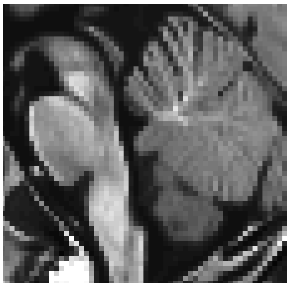 | 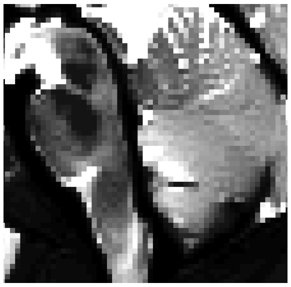 | 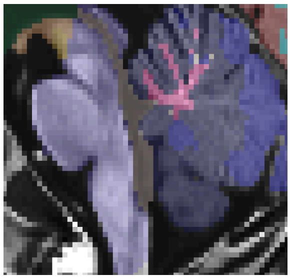 |
| 8mo | 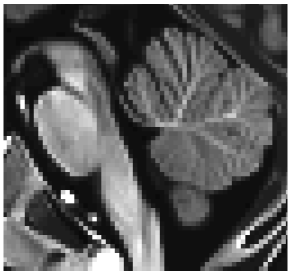 | 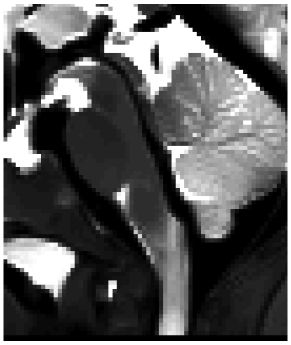 | 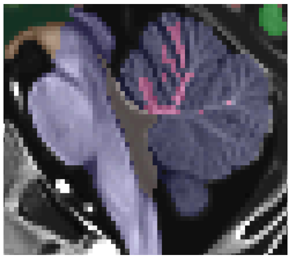 |

Fourth Ventricle Coronal View Figures

| Age | T1 | T2 | Segmentation |
|---|---|---|---|
| 1mo | 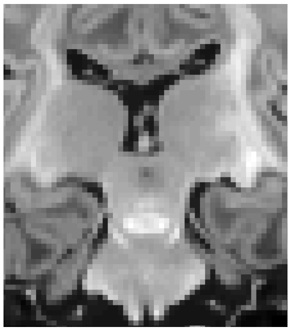 | 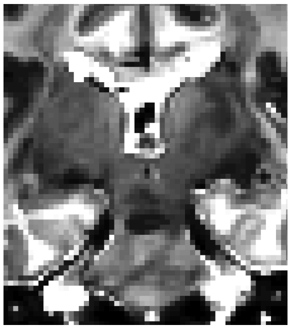 | 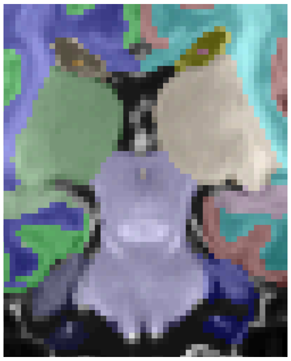 |

| | | | |
|---|---|---|---|
| 4mo | 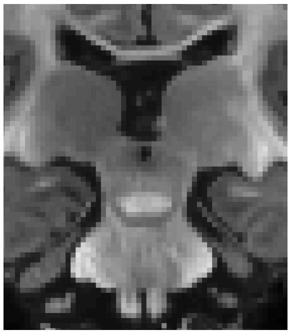 | 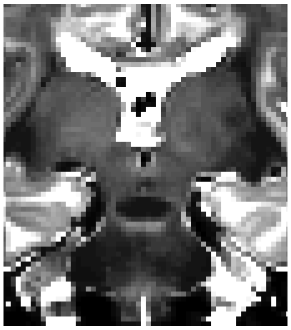 | 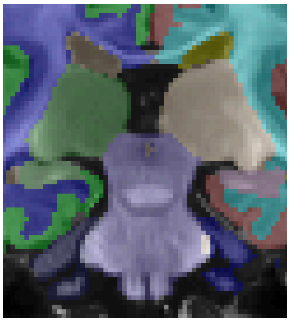 |
| 6mo | 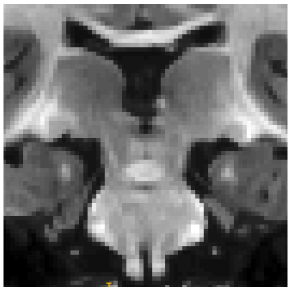 | 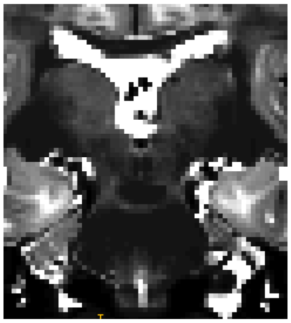 | 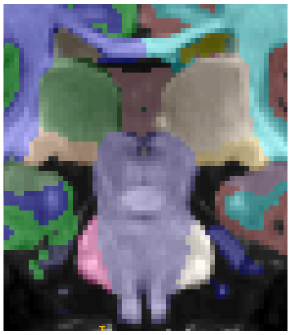 |
| 8mo | 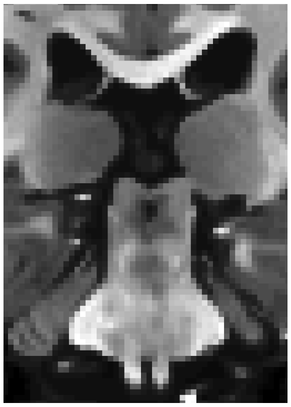 | 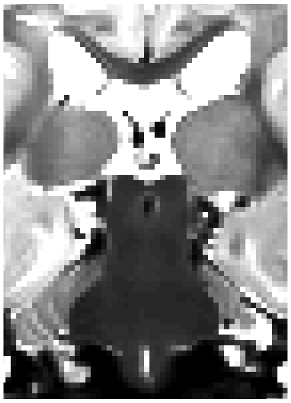 | 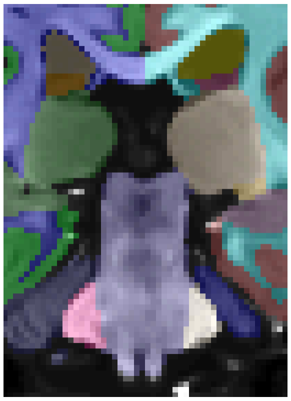 |

### Left/Right Caudate

Based on “The caudate nucleus” section in (Rushmore et al. 2022)

- Defined by the interface with the lateral ventricles, inferiorly by the interface with the thalamus or nucleus accumbens when present, and otherwise by the interface with the adjacent white matter
    - Nucleus accumbens boundary can be a little tricky - the T2 is often helpful.
- See figures to the right for an example of the inferior boundary
- See figures below (T1 left, T2 right) for the posterior boundary (essentially until the caudate disappears):

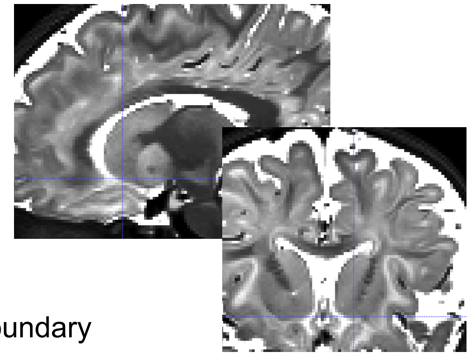

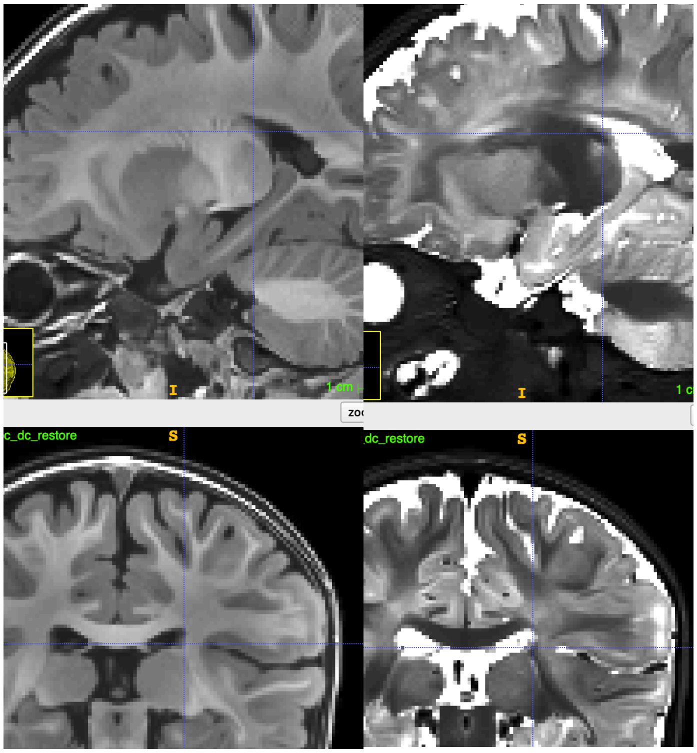


- One frequent issue noted in the caudate is that one hemisphere is over-segmented and the other under-segmented (most commonly observed in 3-4 month old subjects), causing it to be over-extended into white matter (and sometimes even putamen)

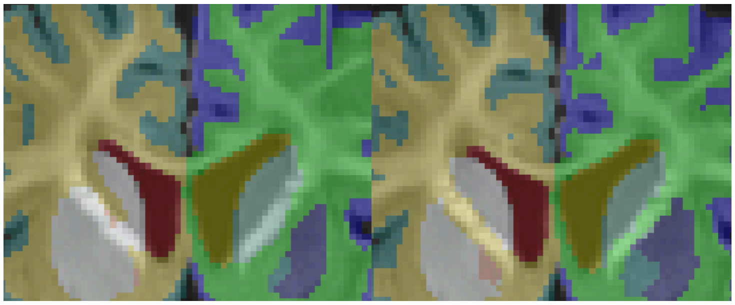

Caudate Figures

| Age | T1 | T2 | Segmentation |
|---|---|---|---|
| 1mo | 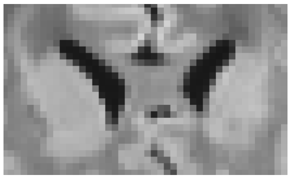 | 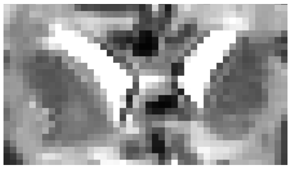 | 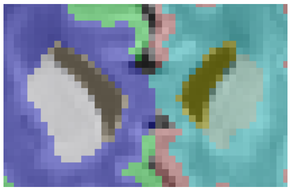 |
| 4mo | 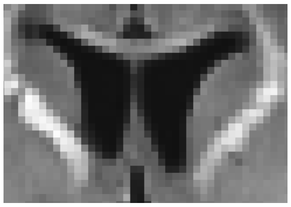 | 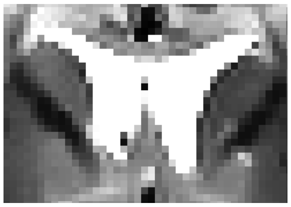 | 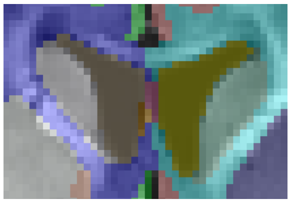 |
| 6mo | 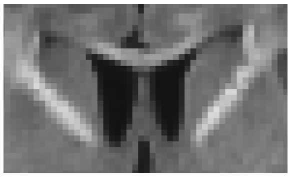 | 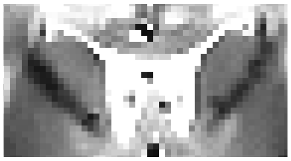 | 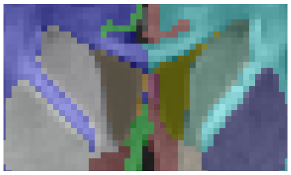 |
| 8mo | 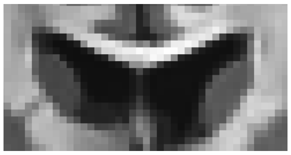 | 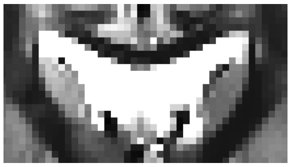 | 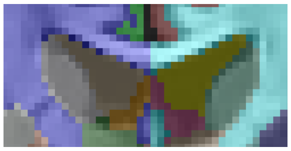 |

### Left/Right Putamen

Based on "The putamen" section in (Rushmore et al. 2022)

- Defined medially by the pallidum (i.e. the putamen is more lateral than the pallidum), laterally by the external capsule, and otherwise by adjacent white matter

- See figure on the left for the boundary with the pallidum (grid line is on the posterior edge bordering the pallidum)

- See figure on the right for the boundary with the external capsule

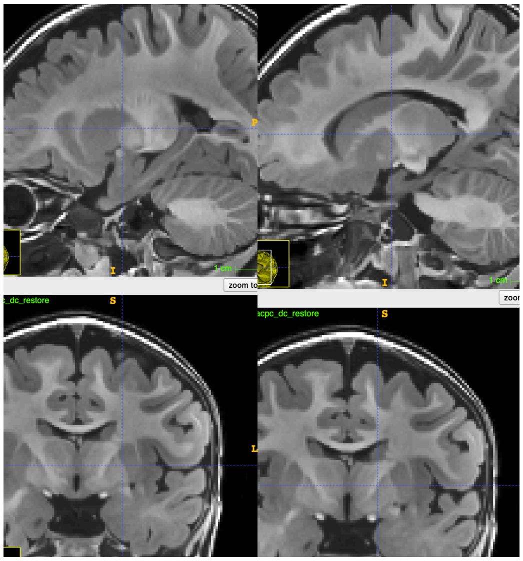

- Common issues in the putamen include both over- and under-segmentation. The screenshot below shows an example of where the putamen was under-segmented (Left: undersegmented, Right: Corrected)

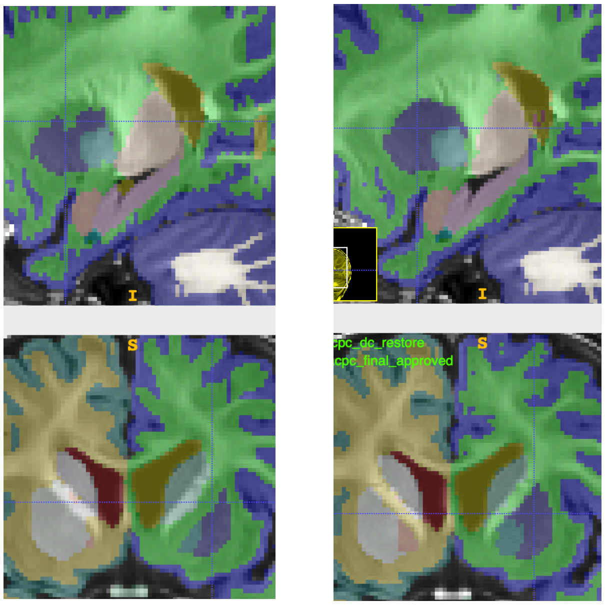

#### Putamen Figures

| Age | T1 | T2 | Segmentation |
|---|---|---|---|
| 1mo | 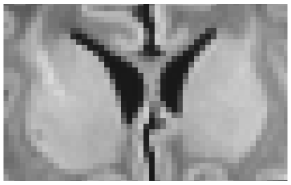 | 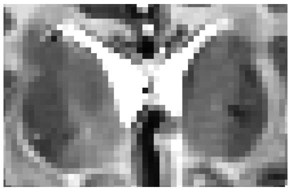 | 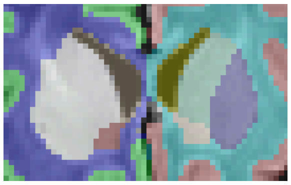 |
| 4mo | 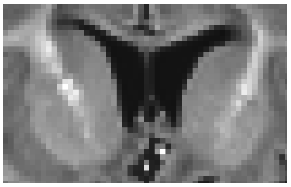 | 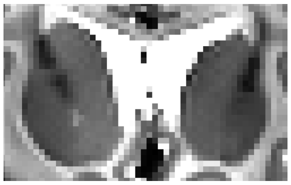 | 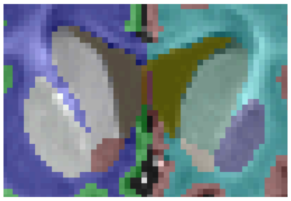 |
| 6mo | 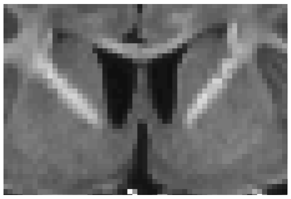 | 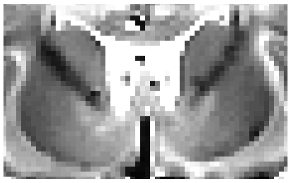 | 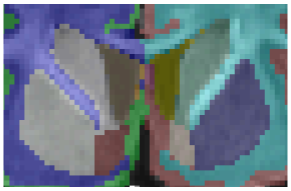 |
| 8mo | 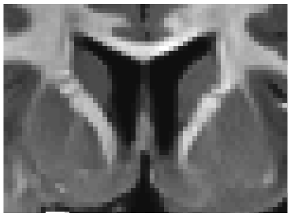 | 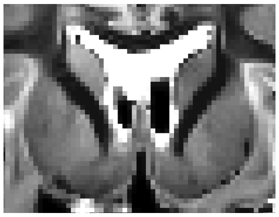 | 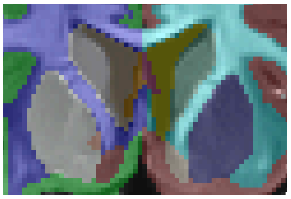 |

### Left/Right Pallidum (Globus Pallidus)

Based on “The globus pallidus” section in (Rushmore et al. 2022)

- Defined by the interface with the internal capsule, inferiorly by the anterior commissure, ansa lenticularis, or nucleus basalis (i.e. things that look like white matter), when present. More medial than the putamen.

- Inferior border where there’s a gap in the white matter (imaginary line between white matter still serves as the boundary - shown in red in the figure on the right)

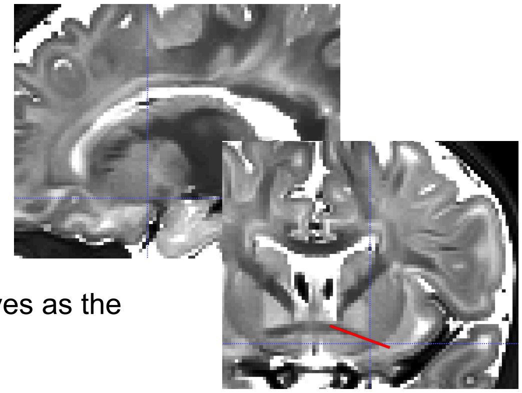

#### Pallidum Figures

| Age | T1 | T2 | Segmentation |
|---|---|---|---|
| 1mo | 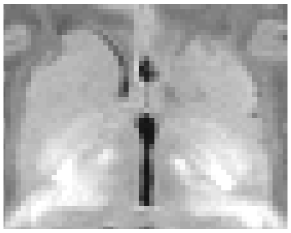 | 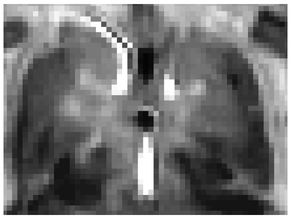 | 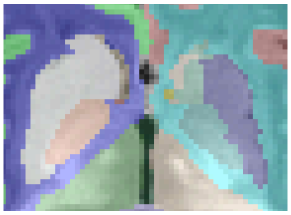 |
| 4mo | 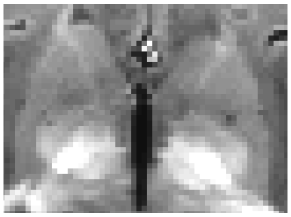 | 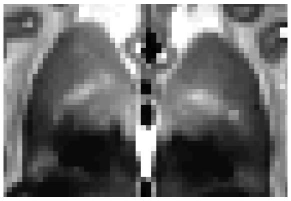 | 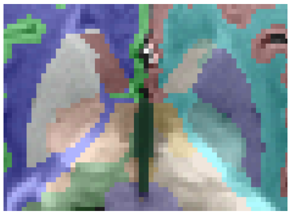 |
| 6mo | 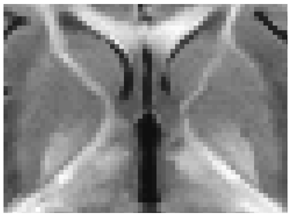 | 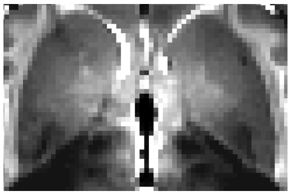 | 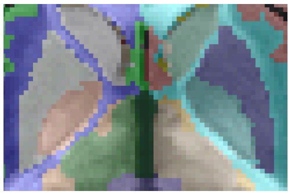 |

| | | | |
|---|---|---|---|
| 8mo | 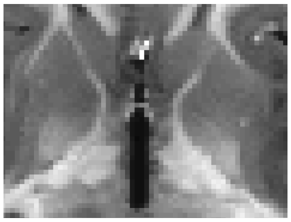 | 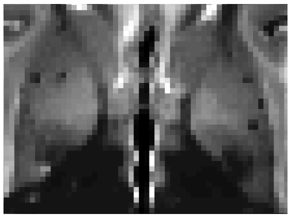 | 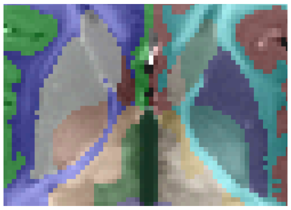 |

### Left/Right Nucleus Accumbens

Based on “The nucleus accumbens” section in (Rushmore et al. 2022)

- Posterior border: appearance of the medial stem of the anterior commissure (gridline is on it in picture to the right)

- The top border in the coronal is an imaginary line drawn between the bottom of the pallidum and the bottom of the putamen. Thus, the nucleus accumbens begins anteriorly when the putamen appears.

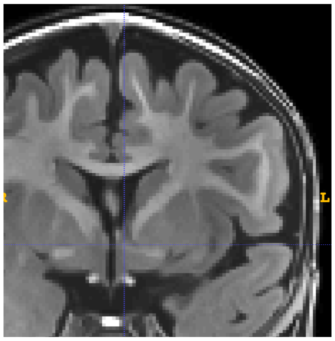

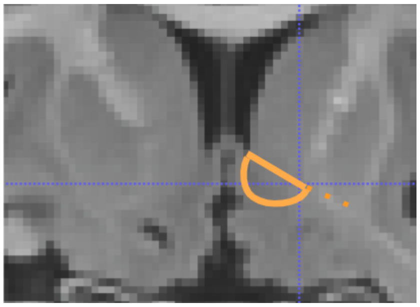

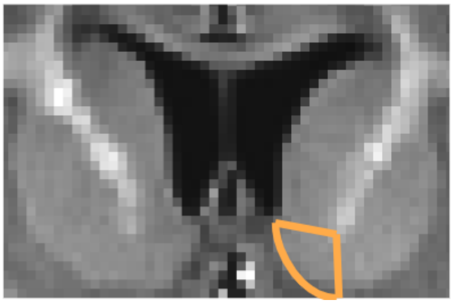

Nucleus Accumbens Figures

| Age | T1 | T2 | Segmentation |
|---|---|---|---|
| 1mo | 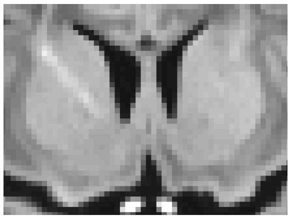 | 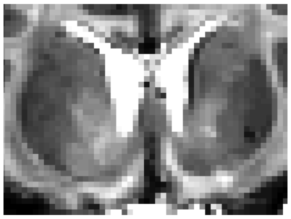 | 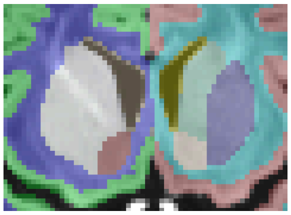 |
| 4mo | 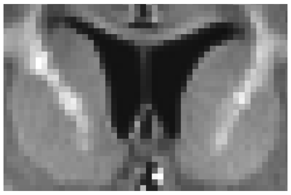 | 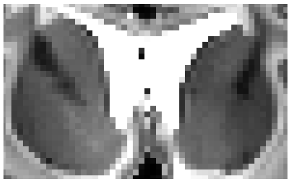 | 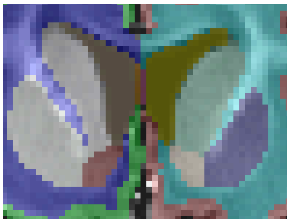 |

| | | | |
|---|---|---|---|
| 6mo | 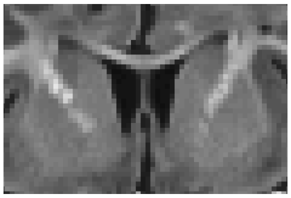 | 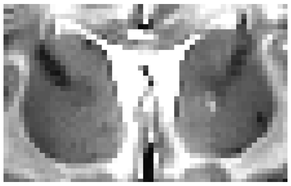 | 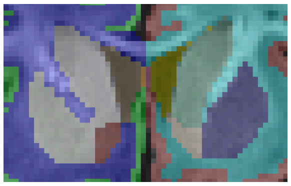 |
| 8mo | 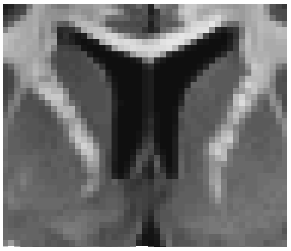 | 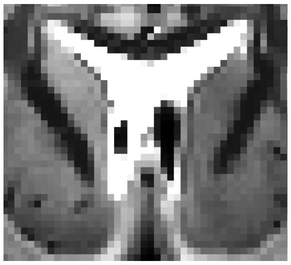 | 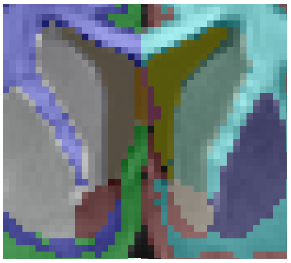 |

### Left/Right Thalamus

Based on “The thalamus” section in (Rushmore et al. 2022)

- The thalamus is a larger structure mostly surrounded by white matter. Grid line in image to the right is on the white matter-thalamus border.

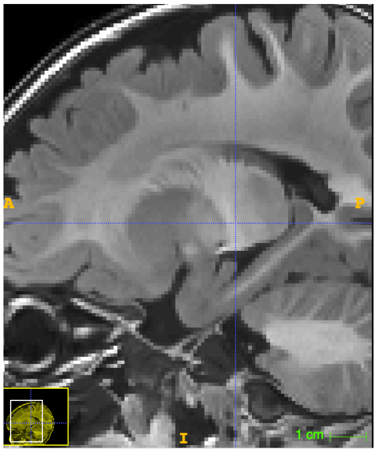


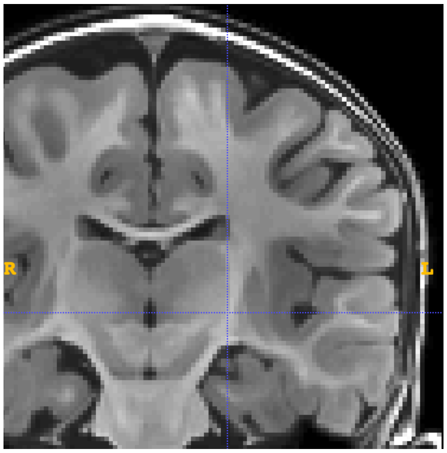


- Inferior border is along the ventral DC (so if you see that labeled, don’t segment past it). The grid line in the picture below is within the ventral DC. The official inferior boundary is the hypothalamic sulcus (which looks like a band of white matter right underneath the thalamus)
    - Known Limitation: The LGN and MGN are included in ventral DC, not in thalamus

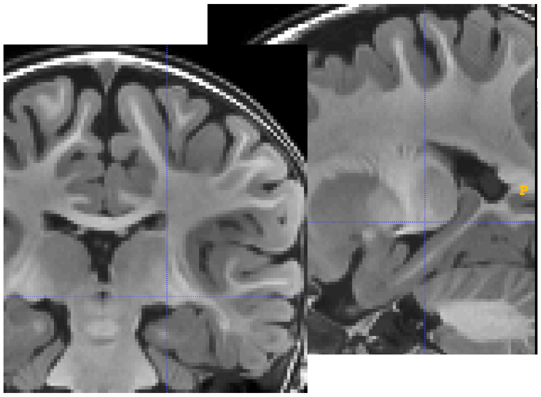


- A common issue is an overlap at the midline in the thalamus - take care that this midline is correct:

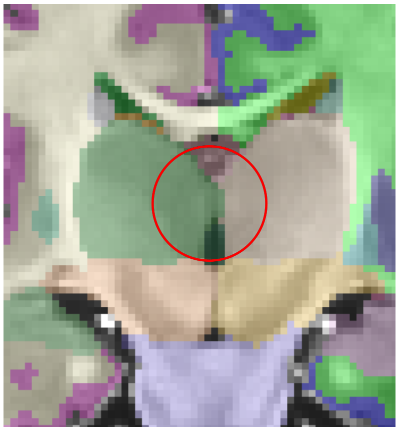

Thalamus Posterior Figures

| Age | T1 | T2 | Segmentation |
|---|---|---|---|
| 1mo | 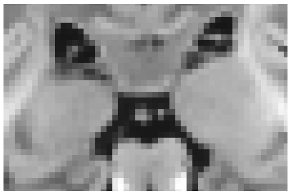 | 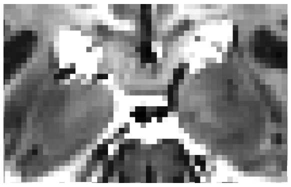 | 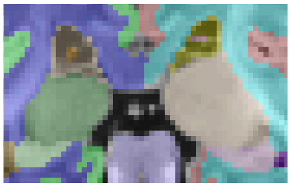 |
| 4mo | 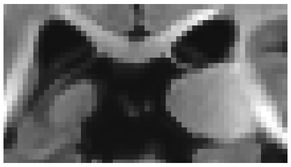 | 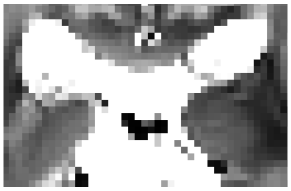 | 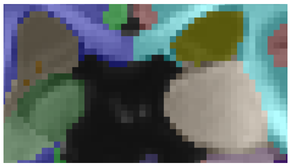 |
| 6mo | 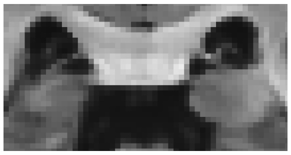 | 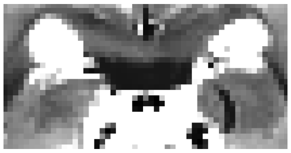 | 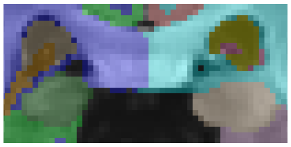 |
| 8mo | 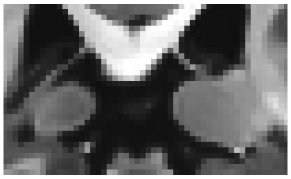 | 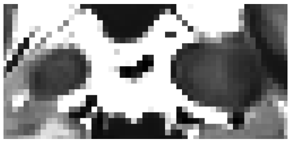 | 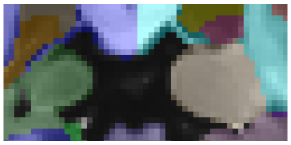 |

Thalamus More Posterior Figures

| Age | T1 | T2 | Segmentation |
|---|---|---|---|
| 1mo | 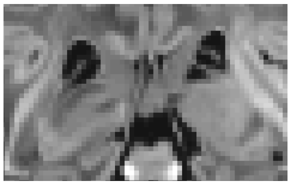 | 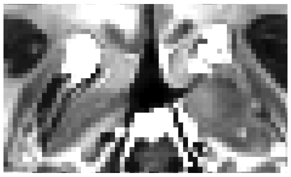 | 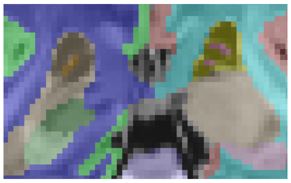 |
| 4mo | 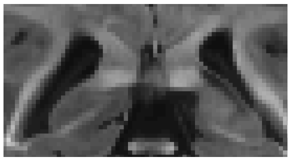 | 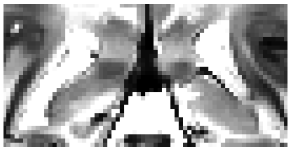 | 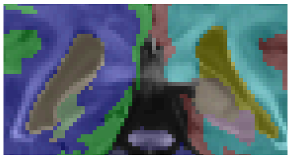 |

| | | | |
|---|---|---|---|
| 6mo | 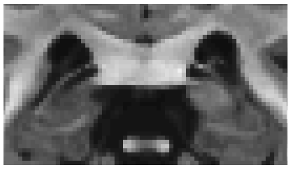 | 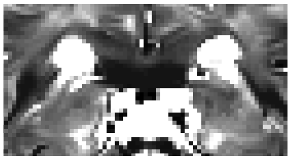 | 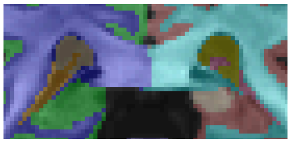 |
| 8mo | 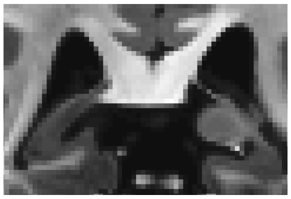 | 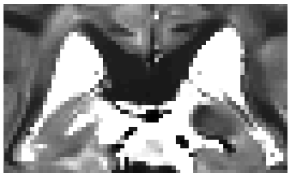 | 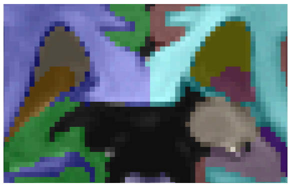 |

Hypothalamic Sulcus

| Age | T1 | T2 | Segmentation |
|---|---|---|---|
| 1mo | 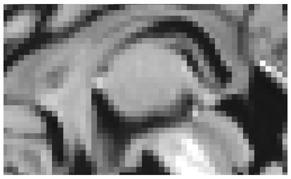 | 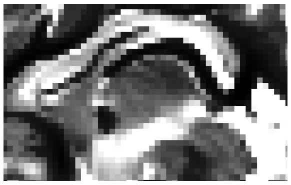 | 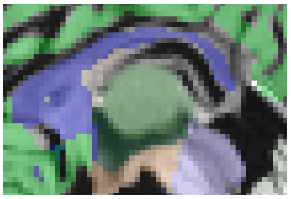 |
| 4mo | 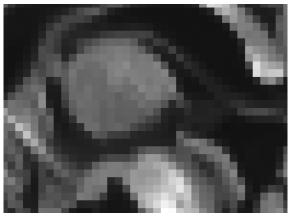 | 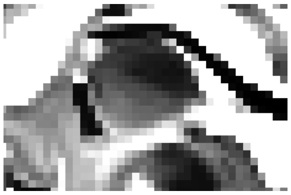 | 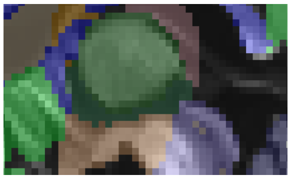 |
| 6mo | 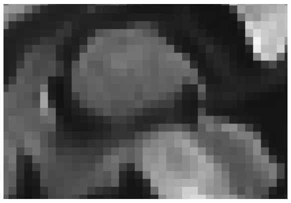 | 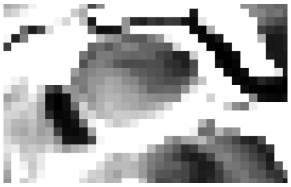 | 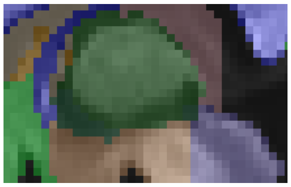 |
| 8mo | 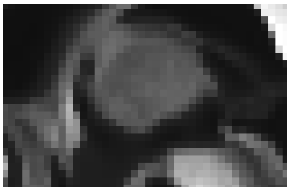 | 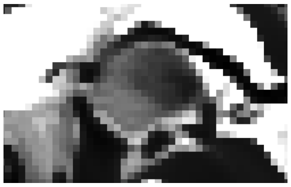 | 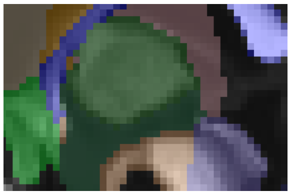 |

### Brainstem

Based on “The fourth ventricle” section in (Rushmore et al. 2022)

- Superior Border (Left) and Inferior Border (Right)

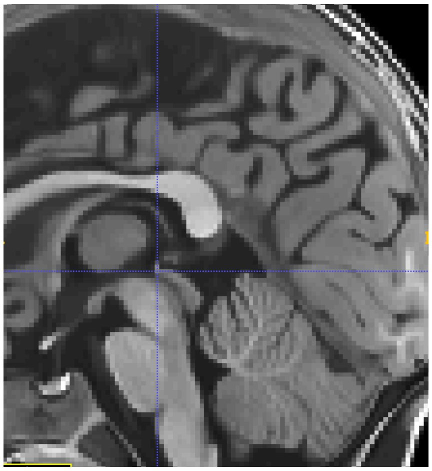

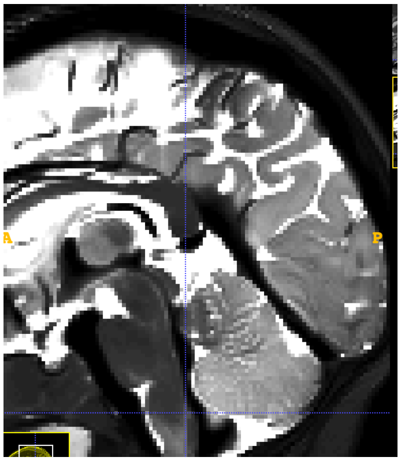

- Use the aqueduct to determine sides (example to the right of a good segmentation)

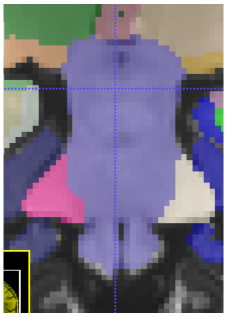

Brainstem Sagittal Figures

| Age | T1 | T2 | Segmentation |
|---|---|---|---|
| 1mo | 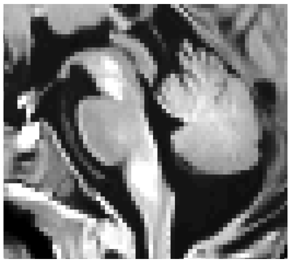 | 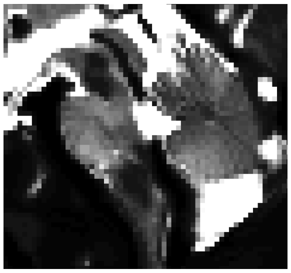 | 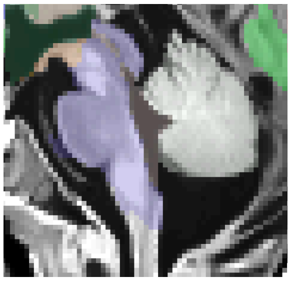 |

| | | | |
|---|---|---|---|
| 4mo | 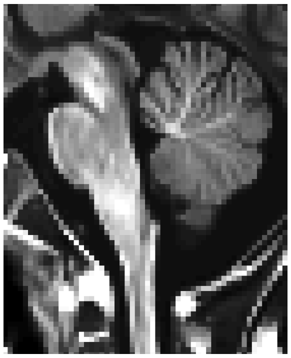 | 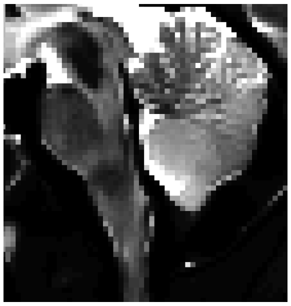 | 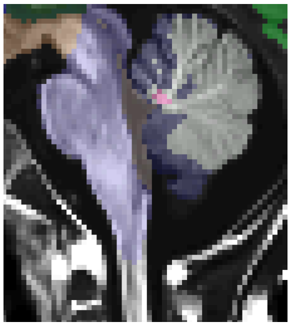 |
| 6mo | 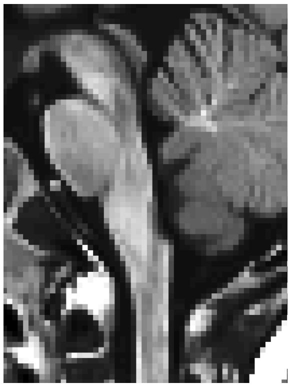 | 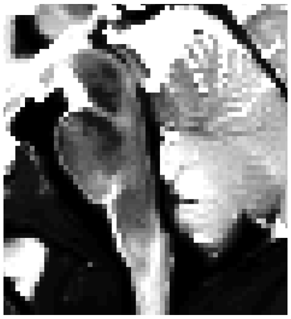 | 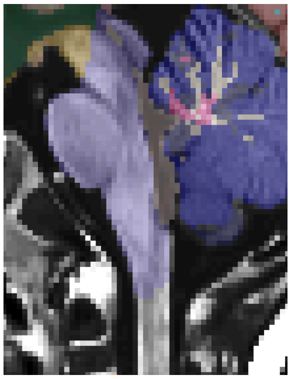 |
| 8mo | 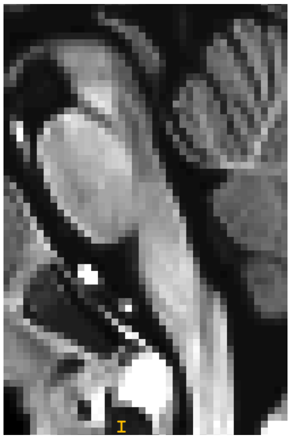 | 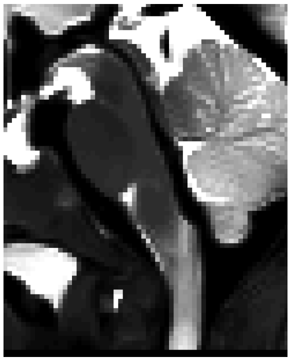 | 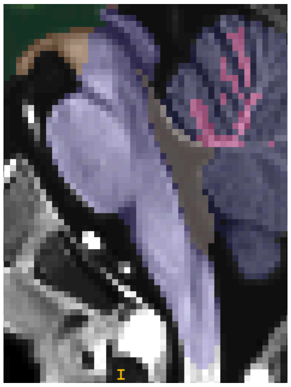 |

Brainstem Posterior Figures

| Age | T1 | T2 | Segmentation |
|---|---|---|---|
| 1mo | 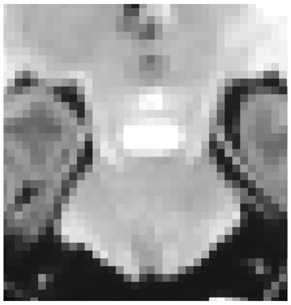 | 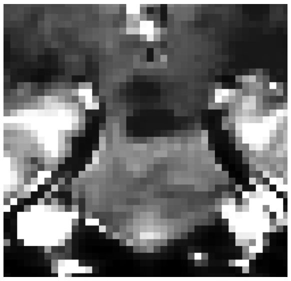 | 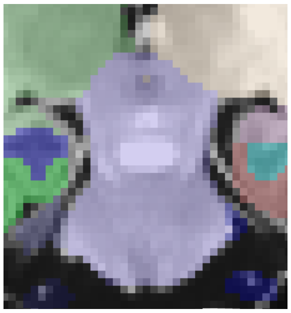 |
| 4mo | 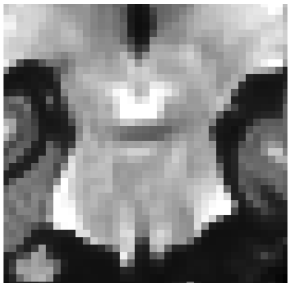 | 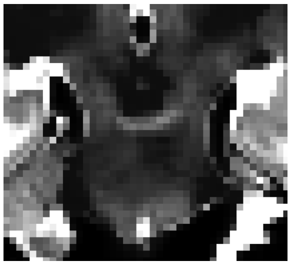 | 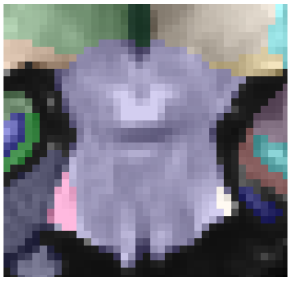 |
| 6mo | 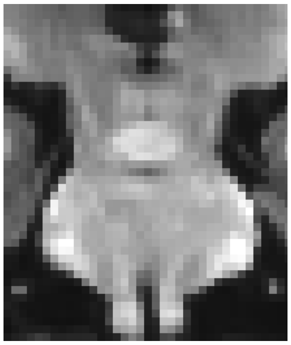 | 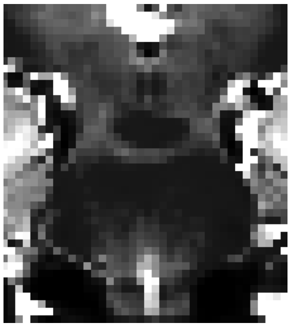 | 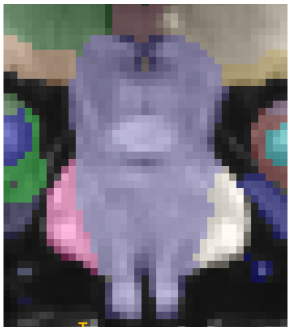 |
| 8mo | 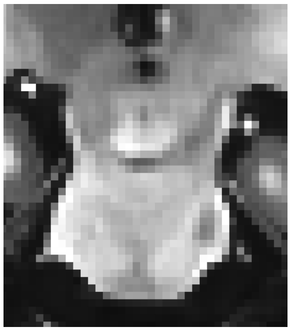 | 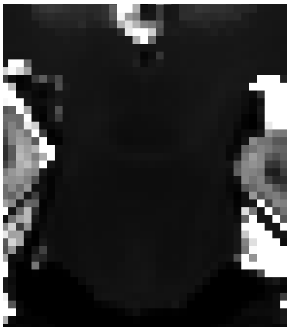 | 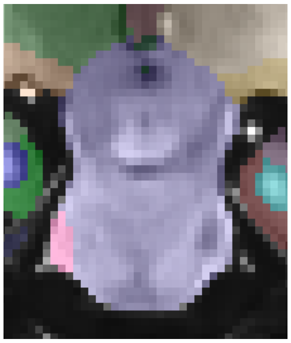 |

### Left/Right Amygdala

Based on (Wedig et al. 2005)

- Anterior (point at which the lateral sulcus closes to form the endorhinal sulcus. L, not closed, R, closed)

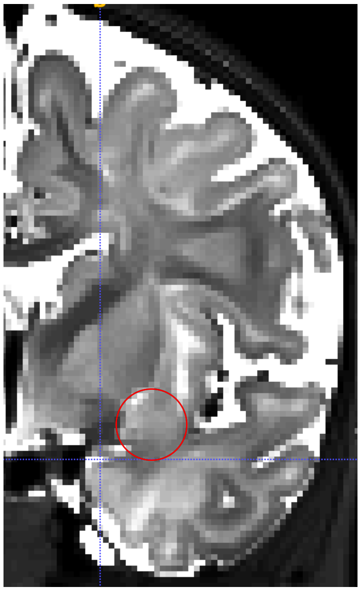

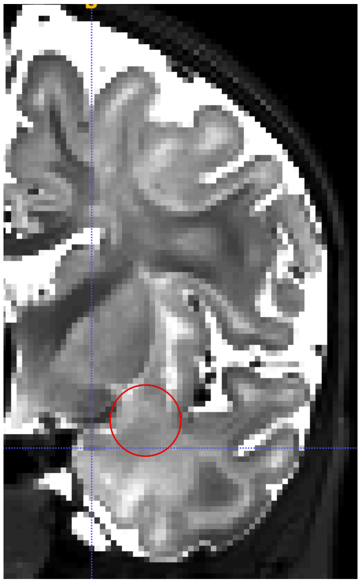

- Sagittal view (below)

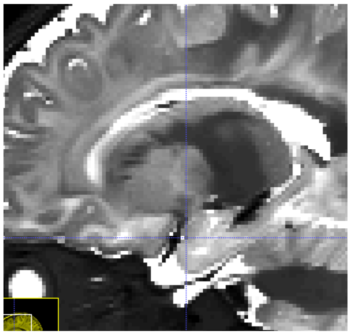

- Posterior: (Mammillary bodies)

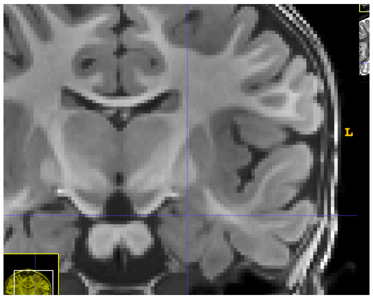

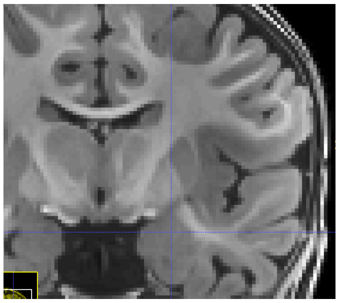

- Medial (line from optic tract (gridline) to tip of white matter in temporal lobe)

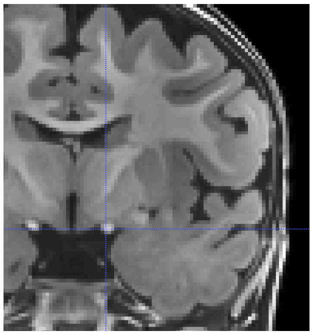

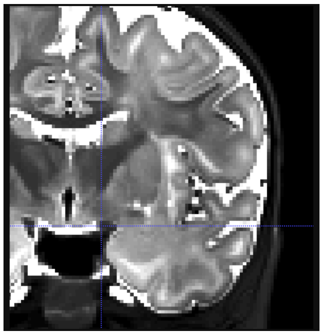

- A common issue in the amygdala is that it is over-segmented into the white matter (Left: Oversegmented; Right: Corrected)

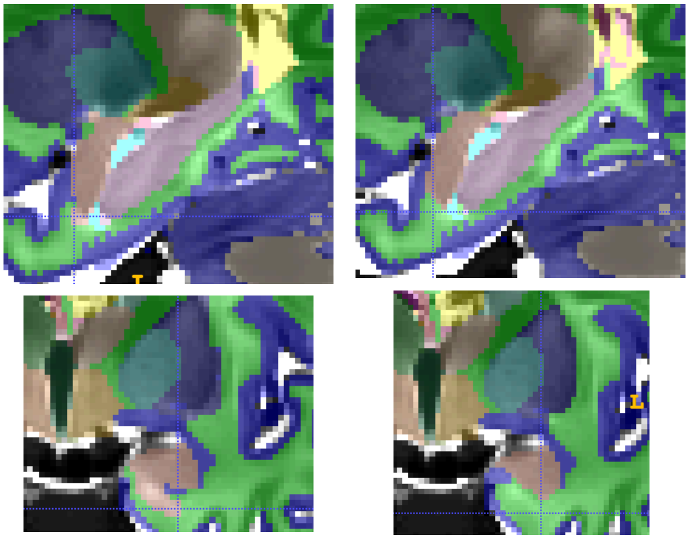

Amygdala Figures

| Age | T1 | T2 | Segmentation |
|---|---|---|---|
| 1mo | 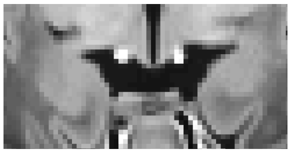 | 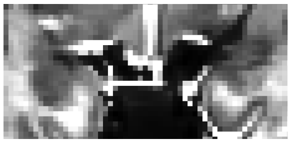 | 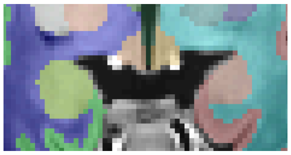 |
| 4mo | 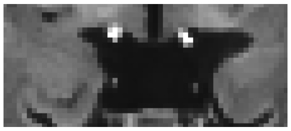 | 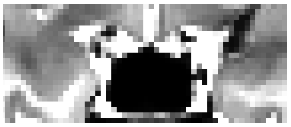 | 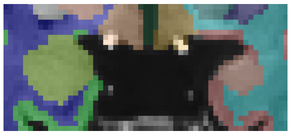 |
| 6mo | 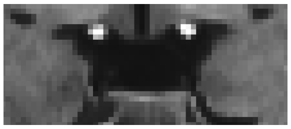 | 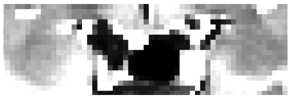 | 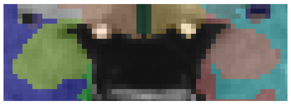 |
| 8mo | 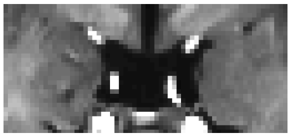 | 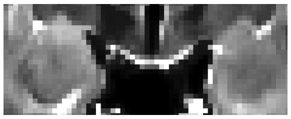 | 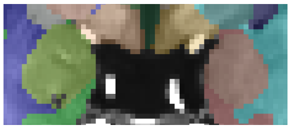 |

### Left/Right Inferior Lateral Ventricle

- As described in the “Lateral Ventricle” section, the lateral ventricle splits from the interior lateral ventricle when the hippocampus appears (when the fornix is disconnected).
    - Depending on the infant, this can be a rather small amount when the ventricle first splits, as seen in some of the examples below
- A common issue is that the inferior lateral ventricle is under-segmented around the amygdala and hippocampus.

Inferior Lateral Ventricle Figures

| Age | T1 | T2 | Segmentation |
|---|---|---|---|
| 1mo | 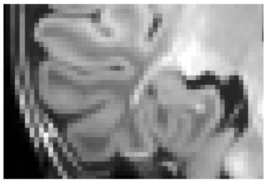 | 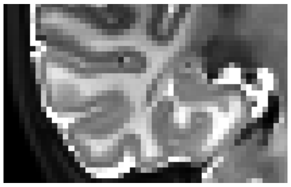 | 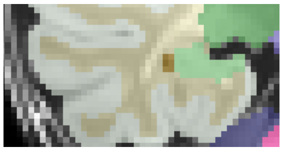 |
| 4mo | 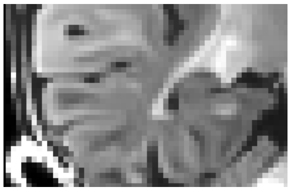 | 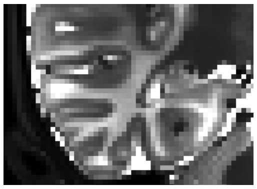 | 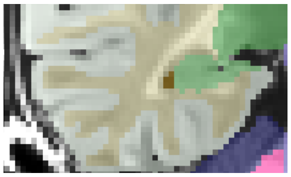 |
| 6mo | 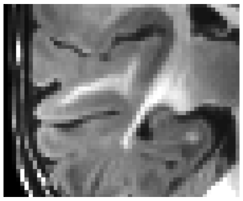 | 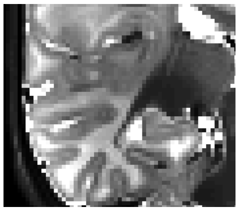 | 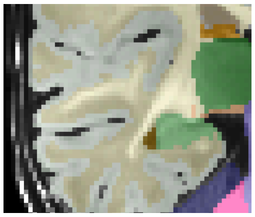 |
| 8mo | 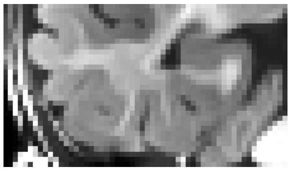 | 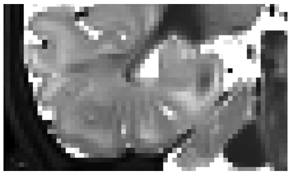 | 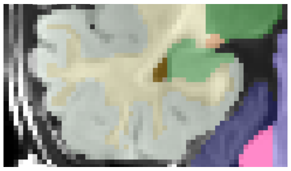 |

### Left/Right Cerebellum

- White matter bounded by cerebellum cortex and brainstem, cortex bounded by CSF and cerebellum white matter
- Known limitations: Quality of white matter edits possible for infants
    - The extension of the cerebellum white matter dentition was near impossible to trace accurately in young infants due to the developing white matter and the thin extensions. Higher resolution scans would be needed to accurately trace the full extensions.
    - Unclear left vs. right hemisphere cerebellum was automatically labeled as “vermis”, a separate label, which was not edited in our protocol

Cerebellum Figures

| Age | T1 | T2 | Segmentation |
|---|---|---|---|
| 1mo | 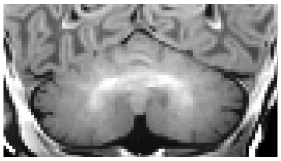 | 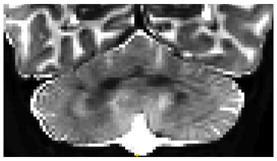 | 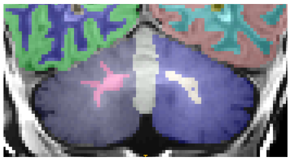 |
| 4mo | 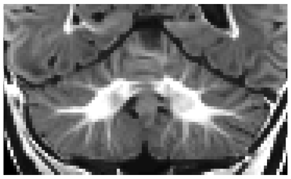 | 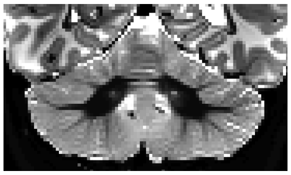 | 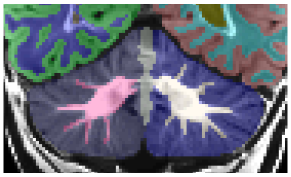 |
| 6mo | 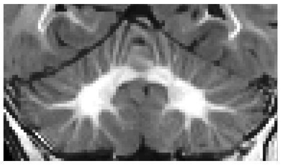 | 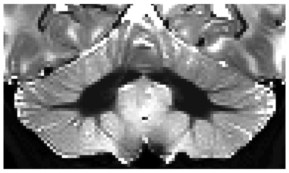 | 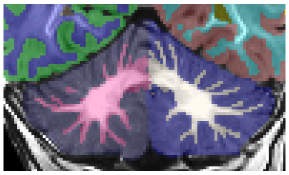 |
| 8mo | 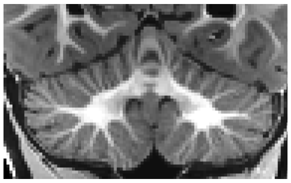 | 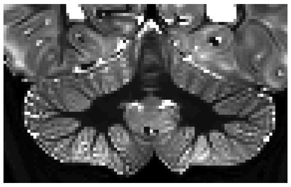 | 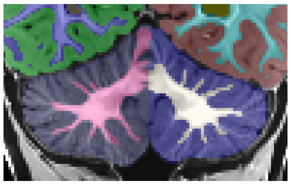 |

### Left/Right Hippocampus

- The hippocampus is bordered by temporal lobe white matter inferiorly and the amygdala anteriorly.
- Posterior end defined as in (Jack 1994, Watson 1992): when fimbria and fornix connect
- 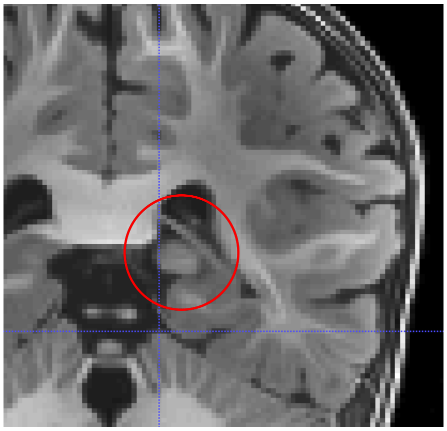 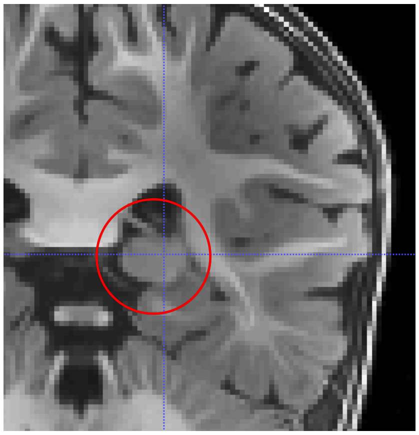
- Known limitation:
  - This definition of the tail ends earlier than the Freesurfer definition. This definition is more consistent with other developmental studies, but will not be consistent with Freesurfer-based segmentations
  - Work is ongoing to add additional hippocampal segmentations to the repository so that researchers can utilize multiple options of hippocampal segmentation types.

##### Hippocampus Figures

| Age | T1 | T2 | Segmentation |
|---|---|---|---|
| 1mo | 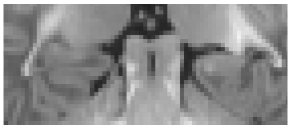 | 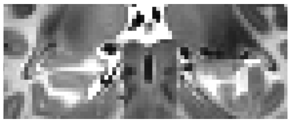 | 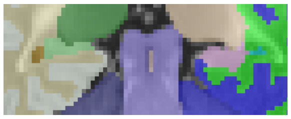 |
| 4mo | 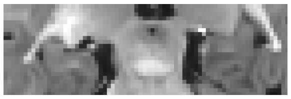 | 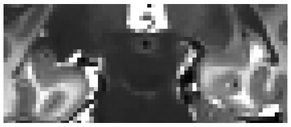 | 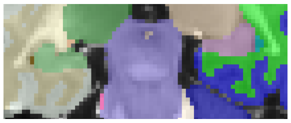 |
| 6mo | 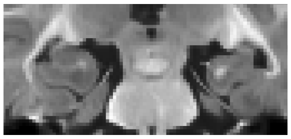 | 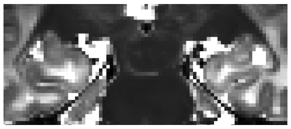 | 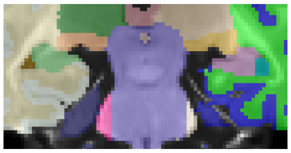 |

| | | | |
|---|---|---|---|
| 8mo | 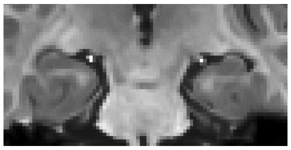 | 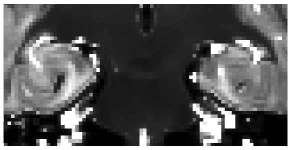 | 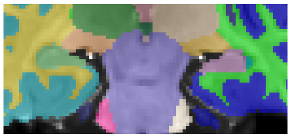 |

Hippocampus Head Figures

| Age | T1 | T2 | Segmentation |
|---|---|---|---|
| 1mo | 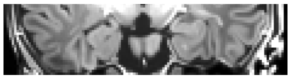 | 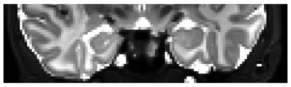 | 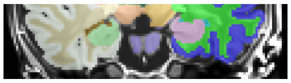 |
| 4mo | 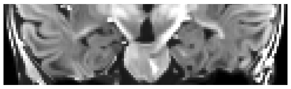 | 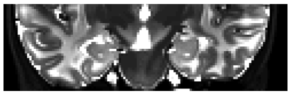 | 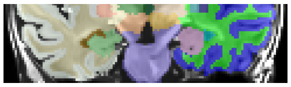 |
| 6mo | 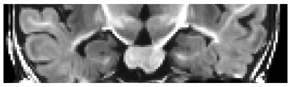 | 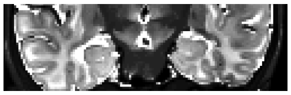 | 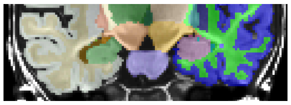 |
| 8mo | 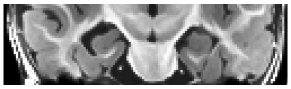 | 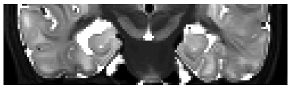 | 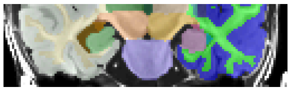 |
